# Characterization of magnetic nanoparticles from the shells of freshwater mussel L fortunei and marine P perna mussels

**DOI:** 10.1101/2022.09.04.506556

**Authors:** Antonio Valadão Cardoso, Clara Carvalho e Souza, Maria Sylvia Dantas, Camila Schults Machado, Erico Tadeu Fraga Freitas, Alisson Carlos Krohling, Veronica A Martins do Rosário, Luiz G Dias Heinene

**Affiliations:** School of Design, State University of Minas Gerais (UEMG); Center For Bioengineering of Invasive Species (CBEIH); Metallurgy Department, School of Engineering (UFMG); Laboratory of Applied Immunology, E Dias Foundation (FUNED); Microscopy Center (UFMG); Center for Nuclear Technology Development (CDTN)

## Abstract

The presence of magnetic nanoparticles in animal species, including humans, has been growing steadily, but none have reported the presence in mollusks apart from the radula of chitons, in 1962. In shells this is the first time. Magnetite (Fe_3_O_4_) nanoparticles were extracted (using three distinct and rather simple protocols) from the shells of freshwater Limnoperna fortunei (Dunker 1857) and marine Perna perna (Linnaeus 1758) mussels and were fully physically-chemically characterized. Due to the spatial distribution, the ferrimagnetic particles in the shells are in low concentration and present a superparamagnetic behavior characteristic of materials of nanometric sizes. Transmission electron microscopy (TEM, especially HRTEM) indicated that the 50-100 nm round magnetic particles are in fact aggregates of 5-10 nm nanoparticles. Using analysis TEM techniques on the shell of the L fortunei we have not found any iron oxide particle at the periostracum layer nor in the calcite layer. Nevertheless, roughly round nanoparticle aggregates of iron hydroxy/oxide were found in the nacar layer, the aragonite layer. Being the aragonite layer responsible for more than 97% of the shell of the L fortunei and considering the estimated size of magnetic nanoparticles we could infer that they might be distributed throughout the nacar layer.

## Introduction

The first and only observation on the presence of magnetite (FeO.Fe_2_O_3_) in mollusks was in the radula of chitons in 1962 [1]. Since then, the presence of biomineralized magnetite in animal species has been growing steadily [2, 3]. But none have reported the presence of iron hydr-oxides in other molluscs apart from chitons and limpets.

We first discovered the presence of magnetic on the shell of the freshwater bivalve golden mussel (*Limnoperna fortunei*) accidently. While in the process of demineralisation of the mussel shell, one of us noticed the presence of black particles aggregates attached to the magnetic stirring bar (PTFE-coated), from a batch of *Limnoperna fortunei* (Dunker 1857) shells (Figures 1.a and 2.a). After repeating the same protocol many times, we were convinced that the extremely small particles come from the shell.

**Fig 1.**
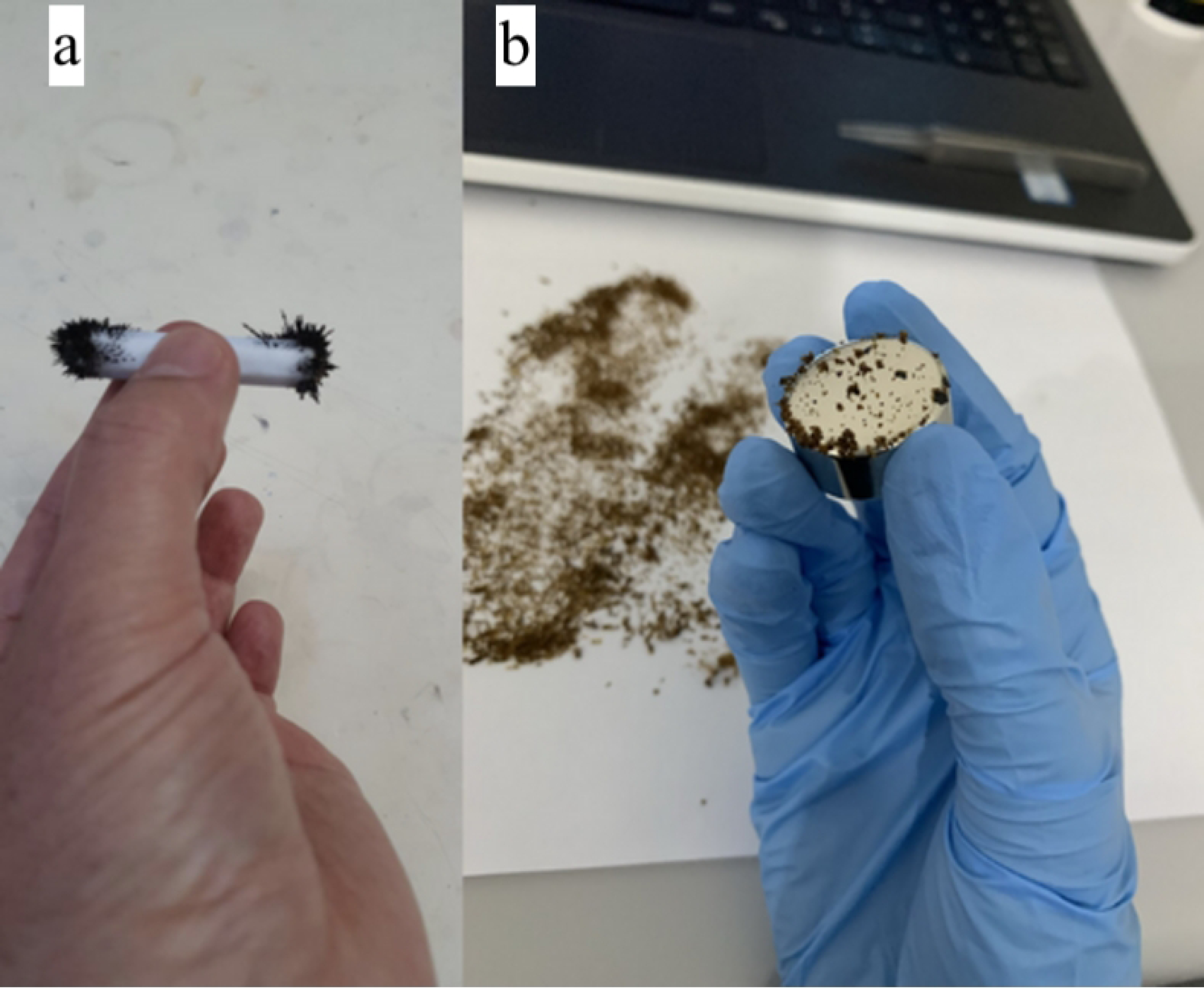
Images of the magnetic particles extracted from the demineralized shells of (a) freshwater bivalve *Limnoperna fortunei* and (b) marine bivalve *Perna perna*. Particles extracted are attached to a magnetic (PTFE-coated) stirrer bar (a) and neodymium (NdFeB) magnet (b)

While verifying in the literature for any clue, we noticed that some scientific papers informed us of the presence of hematite (alpha-Fe_2_O_3_) on the shells of bivalves [4,5,6]. The concentration of hematite on the shell composition is much less than 1% in weight. Because these works aimed at the use of the shell as raw material for plastics, catalysts, etc., they calcined the shell close to 1000 °C. In doing so, the magnetite and any other existing iron compounds have been oxidized and transformed to the higher stable, end phase hematite (Fe_2_O_3_) [7].

The ferrimagnetic magnetite (FeO.Fe_2_O_3_) [8, 9] is an iron ore and has become one of the most interesting crystallographic phases of iron oxide, especially in its nanosized forms. Iron oxide exhibits three different crystalline polymorphs with unique magnetic properties. On heating magnetite, it transforms first into maghemite (gamma-Fe_2_O_3_) at around 300 °C and on hematite (alpha-Fe_2_O_3_) at around 450 °C. These values are for bulk material. Surface to volume ratio is very important and the oxidation of nano magnetite to nano maghemite could occur at temperatures close to ambient.

Further, investigating the literature we noticed that some forty years ago Petit et al [10] had observed the presence of iron-containing micro/nanospheres in the epithelial cells of the mussel mantle.

Historically in 1953 experimental work revealed that one set of valve-mantle, severed apart from the bivalve body and kept in sea water for in vitro studies, could continue to biomineralize [11] for several days. Being an intact system with respect to the relationship between the mantle, the extra-pallial space (ESP) and the shell clearly demonstrates that the calcium was not coming from metabolism. More than seventy years ago experimental work [12] using *valve-mantle preparations* with radioisotope ^45^Ca revealed that calcium present on the mussel mantle remained there and was not supplying the shell biomineralization process. Only in this century advances on the fundamental role of hemocytes on biomineralization have been acquainted and established [13]. The sheet of outer epithelial cells of the mantle, facing the shell, are the frontier where the CaCO_3_ nano-agglomerates (amorphous or crystalline) arrive at the extra-pallial site, the fluid-filled space where CaCO_3_ biomineralization takes place.

In other areas of bivalve research, the capacity of bivalves to produce vesicles with a very complex structure [14, 15] and that acts as precursors for the construction of the byssus has been highlighted.

To continue our investigation on the presence of magnetic on the mussel shells (see Experimental Setup on Figure 2.b) we decided to utilize the same HCl-based demineralization protocol and verify the occurrence of magnetic on the seawater bivalve *Perna perna* (Linnaeus 1758), also known as Brown mussel. As shown in Figure 1.b we observed the presence of magnetic particles (henceforth magnetic powder) attached to a neodymium (NdFeB) magnet after gently moving it above the demineralized shell powder (see video at Supplementary Material). Furthermore, we decided to demineralize the shells using ethylenediaminetetraacetic acid (EDTA), which is a well-known effective quelling organic acid for iron. The aim was to verify the presence of the Fe-EDTA complex using UV-Vis spectroscopy.

**Fig 2.**
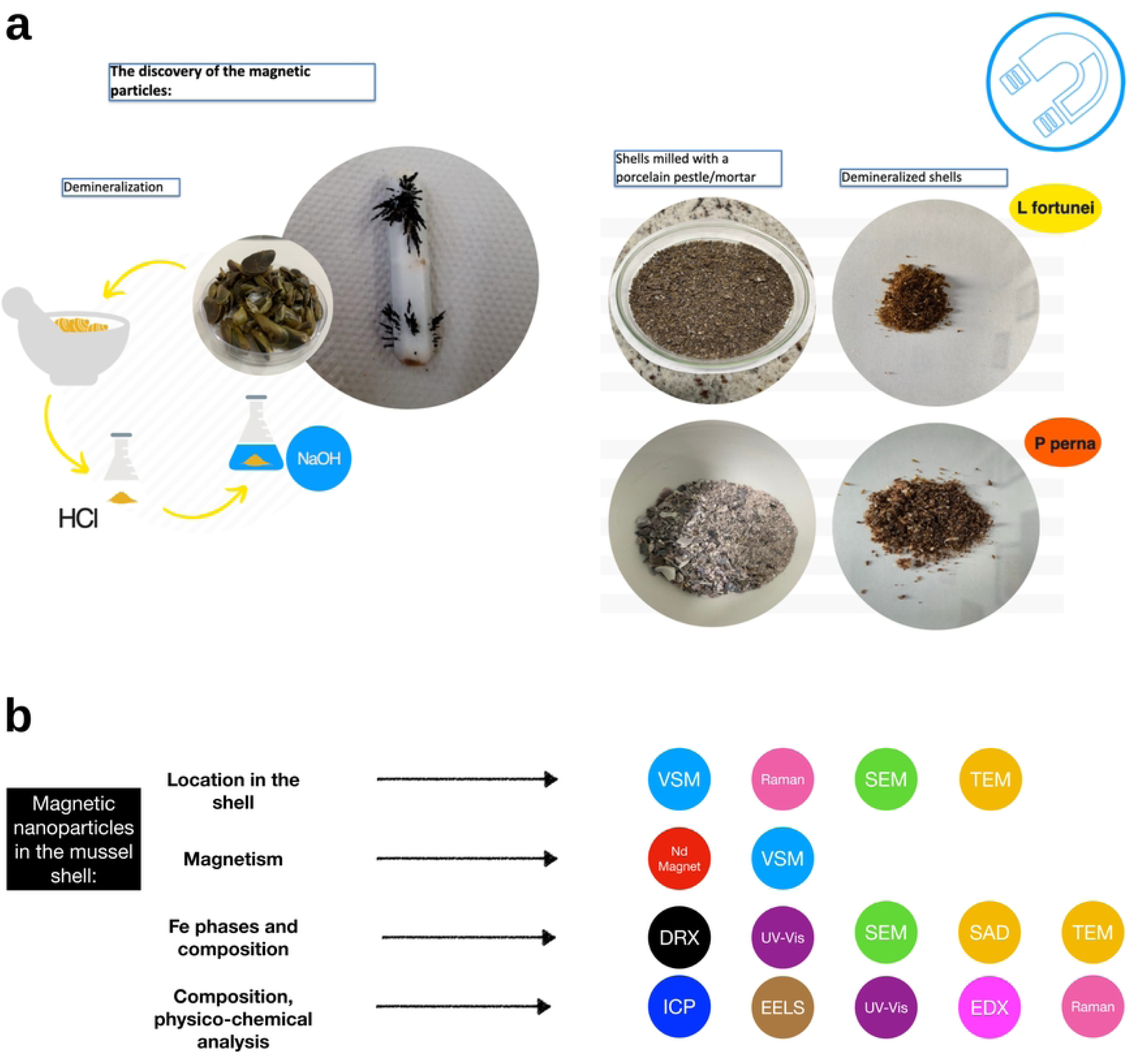
(a) Magnetic particles during demineralization of mussel shells; (b) Experimental setup after magnetic nanoparticles discovery on the mussel shell.

The chemical composition of the *Limnoperna fortunei* was determined using ICP-OES and XRD to quantify the concentration of iron and other metals. To measure the magnetic properties of the shell of the *Limnoperna fortunei* and *Perna perna* we performed magnetometry (Vibrating Sample Magnetometer, VSM). Further, we used scanning electron microscopy (SEM), transmission electron microscopy (TEM), selected area electron diffraction (SAD), energy dispersive X-ray spectroscopy (EDX), and electron energy-loss spectroscopy (EELS) to assess the morphology, the mineralogy, and composition of magnetic powder. We also employed these techniques to analyze the cross-section of the *Limnoperna fortunei* shell in the pursuit to localize the magnetic nanoparticles. Finally, we used Raman spectroscopy to analyze both the magnetic powder and the cross-section of the shell.

In our opinion, the findings may help in advancing our understanding of phase transitions on the interface Biology/Materials Science.

## Materials and Methods

### Specimen collection and preparation

Specimens of *Limnoperna fortunei* were collected at the reservoir of the hydroelectric dam of Volta Grande, latitude 21°46’ South, longitude 42° 32′ West, Minas Gerais, Brazil and kept in an aquarium in our laboratory. Specimens of *Perna perna* were acquired in the local commerce and they came from mussel farms, Santa Catarina Island area, located between latitudes 27°22’ and 27°50’ South and between longitudes 48°25’and 48°35’West, Santa Catarina, Brazil. In the laboratory Perna perna specimens were first kept in the freezer waiting for the demineralization procedure.

### Cleaning, Crushing, Sanding and Powdering shell samples

Detached from the mussels’ body, the shells were first washed in a laboratory sink with running water many times until no trace of organic or inorganic materials (clays, sand, other minerals, etc) were noticed.

The demineralization process was carried out using two different methodologies, to verify the presence of magnetic and iron particles as components of the shells. The golden mussel (L fortunei) shells went through a cleaning process by submersion in a solution of sodium hypochlorite 12% (w/v), to remove all organic contents and other specimens that could be attached to the shells. The 5 minutes incubation at ambient temperature was followed by thorough washing with distilled water and dried in an oven at 70°C. The dried samples were grounded into a fine powder using an analytical mill (IKA A11 basic). This process was the same for both demineralization methodologies used.

### Demineralization with Hydrochloric Acid

The demineralization methodology was adapted [16]. For the process, 1M hydrochloric acid solution was used in the proportion of 40 mL for each 1 gram of shell powder. A total of 250g of powdered shells were added slowly to the acid solution, under constant agitation using a magnetic stirrer. The association of the shell carbonates composition with the acid solution formed bubbles, in this case carbon dioxide, which is evidence of demineralization. After adding all the powder, the mixture was left under constant stirring at room temperature for 6 hours. Subsequently, it was centrifuged at 10,000g for 15 minutes at 25°C, using a Sorvall tabletop centrifuge (model St16-R-Thermo Scientific). The supernatant was discarded and a new solution of 1M hydrochloric acid was added maintaining the proportion 40mL acid/1gram shell followed by an overnight incubation at ambient temperature. The shell powder was washed with ultra-pure water by repeated centrifugation steps, as above, until pH neutrality of the supernatant was observed. After the process, dark colored particles were observed adhered to the magnetic bar, with clear magnetic characteristics. These particles were removed from the magnetic bar and washed, first with distilled water followed by pure ethanol and dried in an oven at 37°C and then weighed.

### Testing Magnetite particles dissolution in 1M HCl solution

Two experiments were performed: one with micrometer size and another with nanometer size magnetic particles. In the first experiment 100 mg of natural magnetite (FeO.Fe_2_O_3_), from Turmalina, Latitude: 17° 16' 59" South, Longitude: 42° 44' 7" West, Minas Gerais, Brazil, particles with 200<x<300 um granulometry range were weighted using an electronic analytical balance, Mettler Toledo, model ME 204/A. After weighting the magnetite sample was immersed in 1M hydrochloric acid (Sigma-Aldrich) solution for 24 hours at 20-25°C ambient temperature. After 24 hs in acid, the magnetite particles were dried in air for many hours at ambient temperature. Finally, the dried sample was carefully collected and then weighted in the same balance. Results of the weighting were recorded.

In a second dissolution test 100mg of 30nm diameter synthetic magnetite particles were immersed in 1M hydrochloric acid solution for 24 hours at 20-25°C ambient temperature. Following the same protocol mentioned above the final weight was recorded.

### Demineralization with Ethylenediaminetetraacetic acid (EDTA)

The use of a second process was adapted [17] from a methodology that focuses on the use of EDTA(Sigma-Aldrich) to demineralize the shell preserving the organic matrix present in it. A solution of 0.014g/L of acid-EDTA was prepared by the gradual addition of the acid to 100 mL of ultra-pure water- (Milli-Q) under constant and vigorous stirring, also, to increase the solubility of the reagent, the temperature was raised up to 50 °C (Solution A). By the end of the process, the pH of the solution was about 3.2, ideal to make a stable binding between EDTA and the iron [18] present in the shell. In 10mL of the initial solution (Solution A), 0.4mg of the shell powder of *Limnoperna fortunei* was added (sample 1). The solution was kept under continuous stirring for 5 days. We observed that the pH was a bit low by the end of the 5th day, getting close to 2.0, for that reason Na-EDTA was added to the solution (0.03 mol/L) for 2 days. By the 7th day, the sample was heated up to 60°C until a yellowish color denoting the presence of iron in solution was observed.

### Direct obtention of magnetic particles without demineralization

Shells of the bivalves *Limnoperna fortunei* and the *Perna perna* (Brown mussel) washed, rinsed and thoroughly cleaned were sanded and powdered using common SiC sandpaper. Then the powdered shell sample was placed on a paper sheet and was scrutinized, from below, with an NdFeB commercial magnet to extract magnetic particles from the powdered shell (See Supplementary Material and Raman experiments). Then the magnetic particles were collected for further experiments and characterization

### ICP-OES

Quantification of the metals present in the shell using ICP-OES was performed in the year 2016 with freshwater L fortunei shell only with an inductively coupled atomic emission spectrometer (ICP OES, Perkin Elmer-Optima 3000). In 2022 we again quantified the metals present in shell samples of the L fortunei and in shell samples of marine P perna mussel, both assays using an ICP-OES (Perkin Elmer DV8300). The CVAAS technique (Cold Vapor Atomic Absorption Spectrometry) was used on both occasions to determine the Hg content. Results expressed as <(value) refer to the quantification limits of the technique. The methodology used was validated according to procedures recommended by official bodies (Clean room class ISO-7 and laminar flow island class ISO-5).

### UV-Vis Spectroscopy

The spectrophotometric measurements were carried out using UV-VIS spectrophotometer (ThermoFisher model Biomate 160) equipped with Peltier thermostatted cell holder, 10mm-pathlength quartz cuvettes and dual silicon photodetector, wavelength range 190–1100 nm. Experiments were carried out with a 1nm wavelength step and solutions two types of solutions were tested: a- Demineralised Limnoperna *fortunei* shells suspensions obtained using aqueous solutions of pure EDTA for demineralization and b- aqueous solutions with pure EDTA.

### Energy Dispersive X-Ray Fluorescence Spectroscopy (EDX)

An EDX Shimadzu equipment, model EDX 7000, were utilized for the experiment and approximately 5 grams of the shell of L fortunei and 5 grams of the shell of P perna were used to analyze de atomic elements concentration (in ppm). Previously the shells of L. fortunei and P. perna were carefully washed with deionized water and brush and dried at ambient temperature.

### X-Ray Diffraction (XRD)

For the analysis, a Multipurpose Diffractometer (model Empyrean, Panalytical) equipped with a copper tube and a two-dimensional PIXEL 2X2 detector was used. The measurement was performed using CuKalpha1 radiation monochromatized with a hybrid double-mirror monochromator. The X-ray diffraction measurement was performed in reflection mode with the sample rotating with a rotation period equal to 4 seconds. Experimental parameters of the measurement were 45 kV tension, 40 mA cathode current and 4-140° angle range.Quantitative analysis of the composition of the samples was performed using the Rietveld method and the quantification of phases was carried out using the Inorganic Crystal Structure Database (ICSD) databases. Shells of L. fortunei and P. perna were carefully washed with deionized water and brush, dried at ambient temperature. Tests were carried out on the powder of the shell, ground with a porcelain pestle and mortar.

### VSM experiments

VSM assays were conducted in a LakeShore Model 7400. Hysteresis measurements were performed with an average of 10 seconds/point; the smallest gap (7.5mm) for the poles of the electromagnet available for the Polychlorotrifluoroethylene (PCTFE) sample holder Kel-F was used. A reference measurement of the Kel-F sample holder was performed under the same conditions, to subtract the sample holder signal from the hysteresis curves of the shells.

The Kel-F sample holder was ultrasonically cleaned in a neutral detergent and isopropyl alcohol before each measurement to remove any contaminants present. In addition, all sample handling, from sample mass quantification to placement in the VSM, was carried out with nitrile gloves and non-magnetic tweezers. We also carried out the process of canceling the magnetic remanence of the magnetometer before each hysteresis curve of the samples.The shells of P *perna* were broken into chunks using a pestle and mortar porcelain. For the experiment with the shells of L *fortunei* a ceramic stylus (ZrO_2_) was used to cut the shell into nine pieces nominated s1, s2,…,s9 (see image of the regions in Figure 14-s1). For both types of shells, the M (magnetization)xH (magnetic field) measurements were normalized by the respective masses of each individual shell piece.

In a third VSM experiment L *fortunei* powdered shell sample was placed on a paper sheet and was scrutinized, from below, with an NdFeB commercial magnet to extract magnetic particles from the powdered shell (See S1 video in Supplementary Information). Then the magnetic particles was placed on the VSM equipment following the same procedure described above.

### TEM e SEM Experiments

Transmission electron microscopy (TEM) and scanning electron microscopy (SEM) were performed at the Center of Microscopy-Universidade Federal de Minas Gerais, Brazil (www.microscopia.ufmg.br). The experiments were performed with the magnetic powder extracted from the shells and a cross-section of the *Limnoperna fortunei* shell. For the SEM, a small amount of the magnetic powder sample was placed on the carbon tape attached on an aluminum stub. The excess powder was removed from the carbon tape. For the TEM, a small amount of the magnetic powder was dispersed in a 1.5 mL Eppendorf tube filled with distilled water. It was manually agitated and sonicated for 5 min in an ultrasound bath. Then, a drop of the supernatant was placed in a holey carbon-coated Cu-TEM grid of 300 mesh. The cross-section of the *Limnoperna fortunei* shell was cut with an ultramicrotome (Leica UC6) using a diamond knife (Diatome Utra 35). The surface of the shell portion cut was analyzed under the SEM and thin section (100 nm thick) under the TEM.

For TEM experiments, we used a thermionic LaB6 FEI Tecnai G2-20 SuperTwin microscope, operated at 200 kV, coupled with a Gatan Image Filter (GIF) Quantum SE, and an energy dispersive X-ray spectrometer, silicon drift detector Oxford TEM Xplore, with an 80mm² window. Electron energy-loss spectroscopy (EELS) and energy dispersive X-ray spectroscopy (EDX) were performed in scanning TEM (STEM) mode. The crystallographic analysis was done with selected area electron diffraction (SAD). Local elemental analyses were carried out by EELS and EDX, and phase analysis was also done by EELS. As for the SEM experiments, we used a field emission gun FEI Quanta 3D microscope, coupled with a Bruker SDD-EDX.

### Raman spectroscopy

For Raman analysis, a LabRam-HR 800 spectrometer (Horiba Jobin Yvon) equipped with a He-Ne laser (excitation at 632.8 nm) and an Olympus BX41 microscope with lenses of 10X, 50X, 100X and 100XLWD were used. The laser was focused at 1-2 um^2^ area of the sample (with the 100X lens). Scattered light is collected by a monochromator and detected by a liquid nitrogen cooled CCD. The spectrum scans from 100 to 4200 cm^-1^ with a 1.1 cm^-1^ pitch. For most of our experiment we scanned an 80-1250 cm^-1^ window, acquisition time 30s and was acquired 10 times to increase the signal to noise ratio. To avoid local heating in the sample, which can result in thermal transformation, the power of the laser was tested.

## Results

### Testing Magnetite particles dissolution in 1M HCl solution

A careful test was conducted to verify if the 1M HCl solution utilized for the demineralization of the bivalve shell could dissolve the iron present on the shell and re-precipitate as magnetite nanoparticles. The magnetite particles could therefore be an artifact from the HCl shell dissolution experiment. However, after 24 hrs in 1M HCl the same initial weight (100 mg of magnetite particles) was recorded indicating that no magnetite mass dissolution occurred in this period.

However, in the second dissolution test with 100mg of 30nm magnetite particles nearly all the magnetite nanoparticles have been dissolved after 24 hours in 1M HCl. Just 1.5 g was measured as the final weight.

### Direct obtention of magnetic particles without demineralization

Due to the possible influence of the biomineralization process with HCl 1M on the appearance of magnetic nanoparticles we decided to simply grind cleaned and dried shells with commercial sandpaper (SiC) and verify the occurrence of magnetic particles.

By sanding the shell of *L fortunei* with commercial sandpaper we were able to collect magnetic samples from the powder with the aid of a NdFeB magnet (Supporting Information, S1- Video: Sanded L fortunei mussel shell: collecting magnetic particles). Further we measured the saturation magnetization Ms (See VSM results) of the magnetic particles collected with the of a magnet.

### Chemical shell composition of Limnoperna fortunei using ICP – OES and EDX

Table 1 shows the concentrations of different metals present on the freshwater Limnoperna fortunei mussel shell. Obviously, the concentration of calcium is very high considering that is the main component of the shell. Sr, Al, Fe, Mn, Mg and Na are the only metals that achieved concentrations of a fractions of percentage. Clearly these metals and other heavy metal traces were in the freshwater and were assimilated at or beyond the gills. The highest is Na which achieves a decimal percentage probably because sodium is essential in all metabolic processes. The other metal elements concentrations are very close to the values obtained elsewhere [19], especially for calcium ions which as CaCO_3_ constitutes more than 90% of the shell composition. The iron concentration in the shell of P perna is close to the one obtained in other studies of marine bivalves [20, 21] but the iron concentration in the shell of L fortunei differs from the iron composition in the shell of other freshwater bivalves.

**Table 1.**
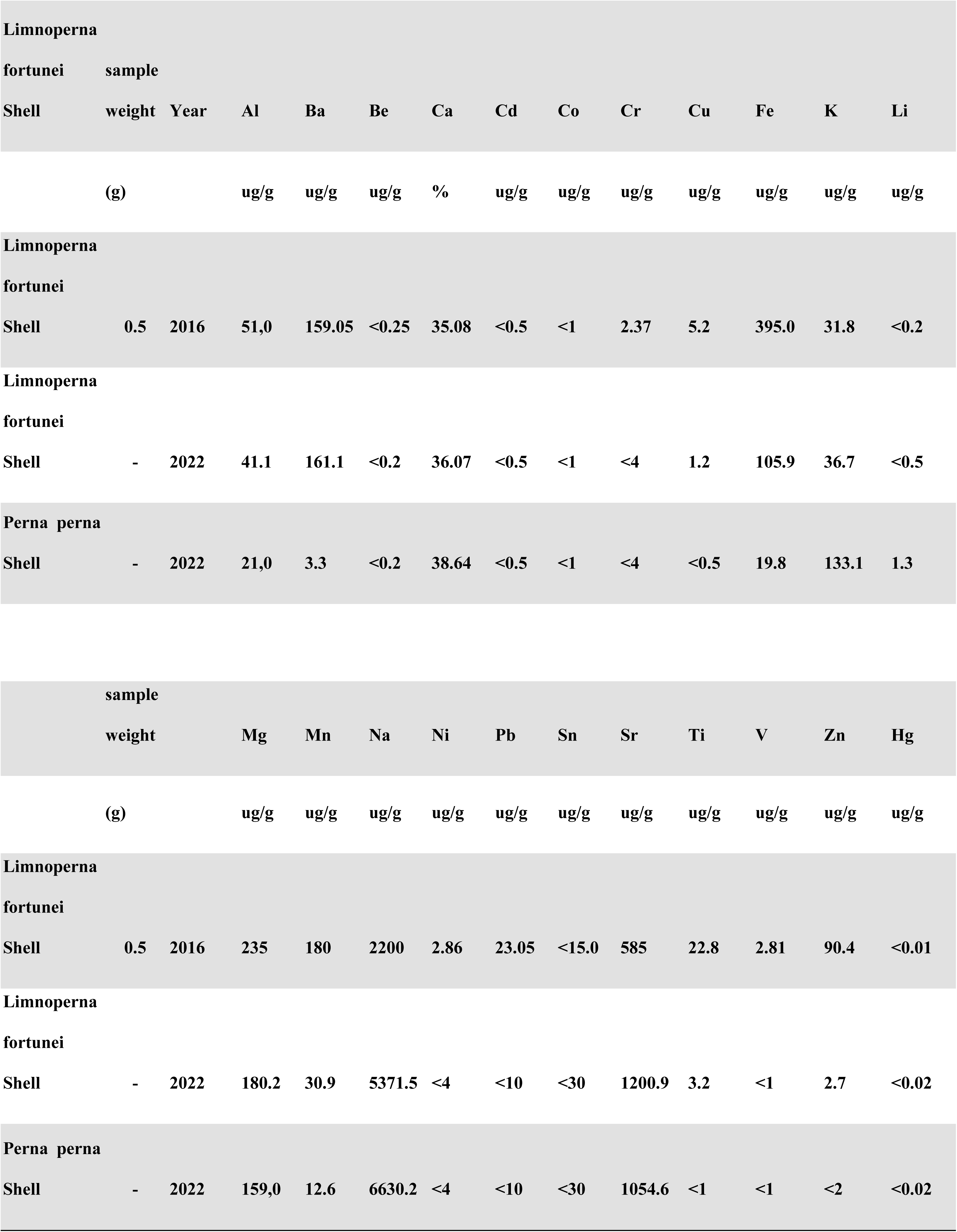
Metal element composition of the shell of the Limnoperna fortunei shell using ICP-OES technique(in mg/g and %).

The results of EDX are presented in the Supplementary Information (S3 Table). They are in line with the results obtained with ICP-OES and indicate that the concentration of iron on the L *fortunei* shell is higher than in P *perna* shell.

### UV-Vis Spectroscopy

Figure 3 presents the results of UV spectral absorption of the magnetic particles extracted from the shell of *L fortunei* in Na-EDTA solutions and demineralized shell liquor in EDTA and Na-EDTA. NaFe(III)EDTA only, Na-EDTA only, and pure EDTA solutions are present for comparison. As it is shown EDTA does not absorb in the 190-450nm region while the solution with the demineralized *Limnoperna fortunei* shell in EDTA solution (see solution preparation in Methods) shows absorption in the 250-260 nm indicating the formation of EDTA-Fe complex. The UV absorption of the magnetic particles in Na-EDTA solution and NaFe(III)EDTA solution both peaked at 257 nm. At the lowest pH value higher dissolution rates are realized as protons reach the activated state for the breaking of the residual O-Fe bonds [22].

**Fig 3.**
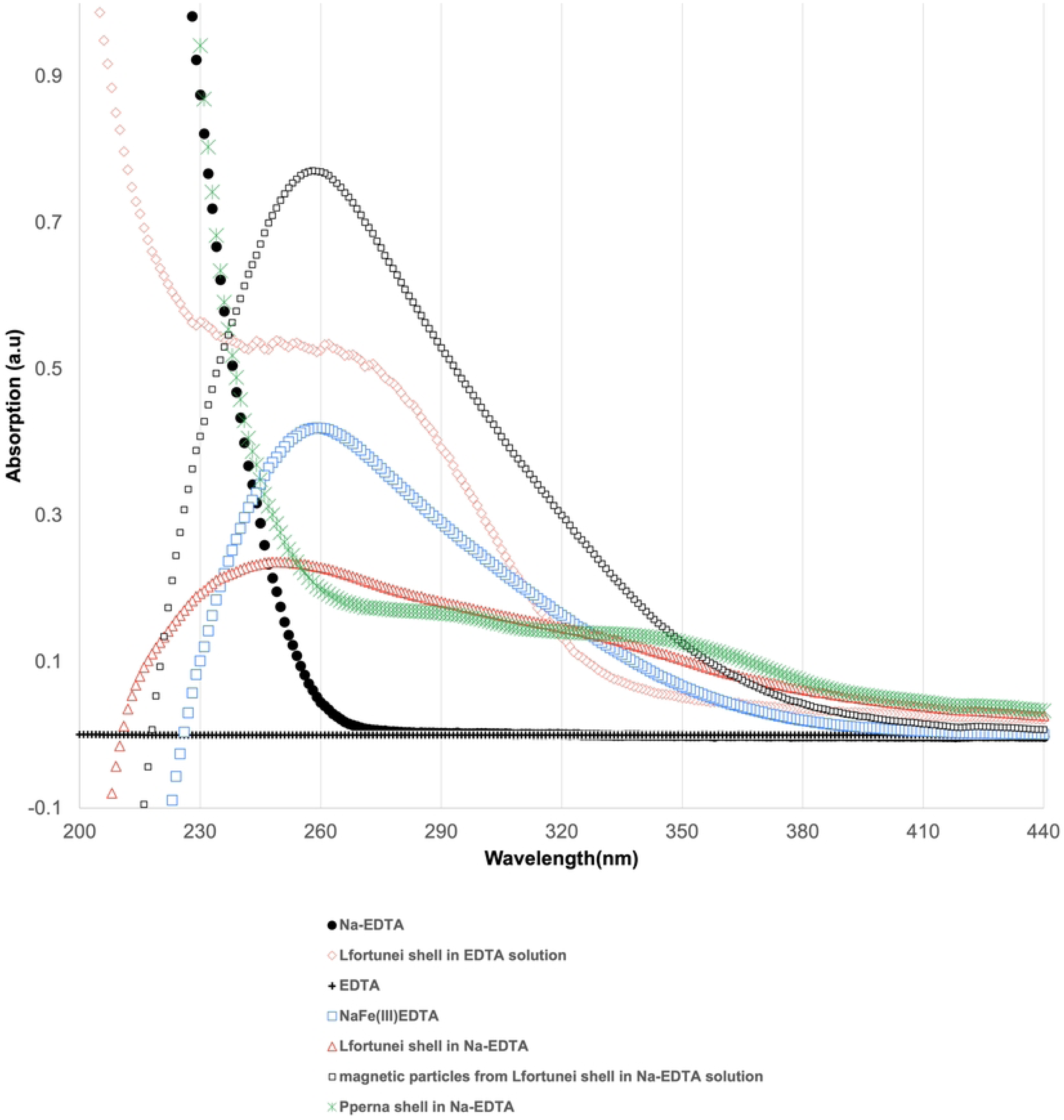
UV-Vis spectra of ethylenediaminetetraacetic acid (EDTA)(dash), Na-EDTA (circle), NaFe(III)EDTA(square) in solution, Golden mussel (Limnoperna fortunei) shell powder in EDTA(diamond) and in Na-EDTA solution(triangle) and magnetic particles extracted from L Fortunei shell in Na-EDTA. Magnetic particles in Na-EDTA and NaFe(III)EDTA have absorptions at 257 nm; EDTA does not absorb in the UV part of the electromagnetic spectrum while demineralized L fortunei shell in Na-EDTA have absorption peak in the region of 230-260 nm.

### X-Ray Diffraction

Figure 4.a and 4.b presents the x-ray diffractograms of the shell, respectively, of the *Limnoperna fortunei* and of the *Perna perna* mussels. Aragonite is the main phase present on the shell of the *L fortunei* mussel and on the Perna *perna* mussel was the only phase detected by DRX. Table 2 shows the concentration of aragonite and calcite phases indicating that the *L fortunei* shell is nearly all composed of aragonite being the calcite, a very thin layer attached to the periostracum [23]. The presence of the iron phases was not detected by XRD as these phases occur in concentrations lower than the detection limits.

**Fig 4.**
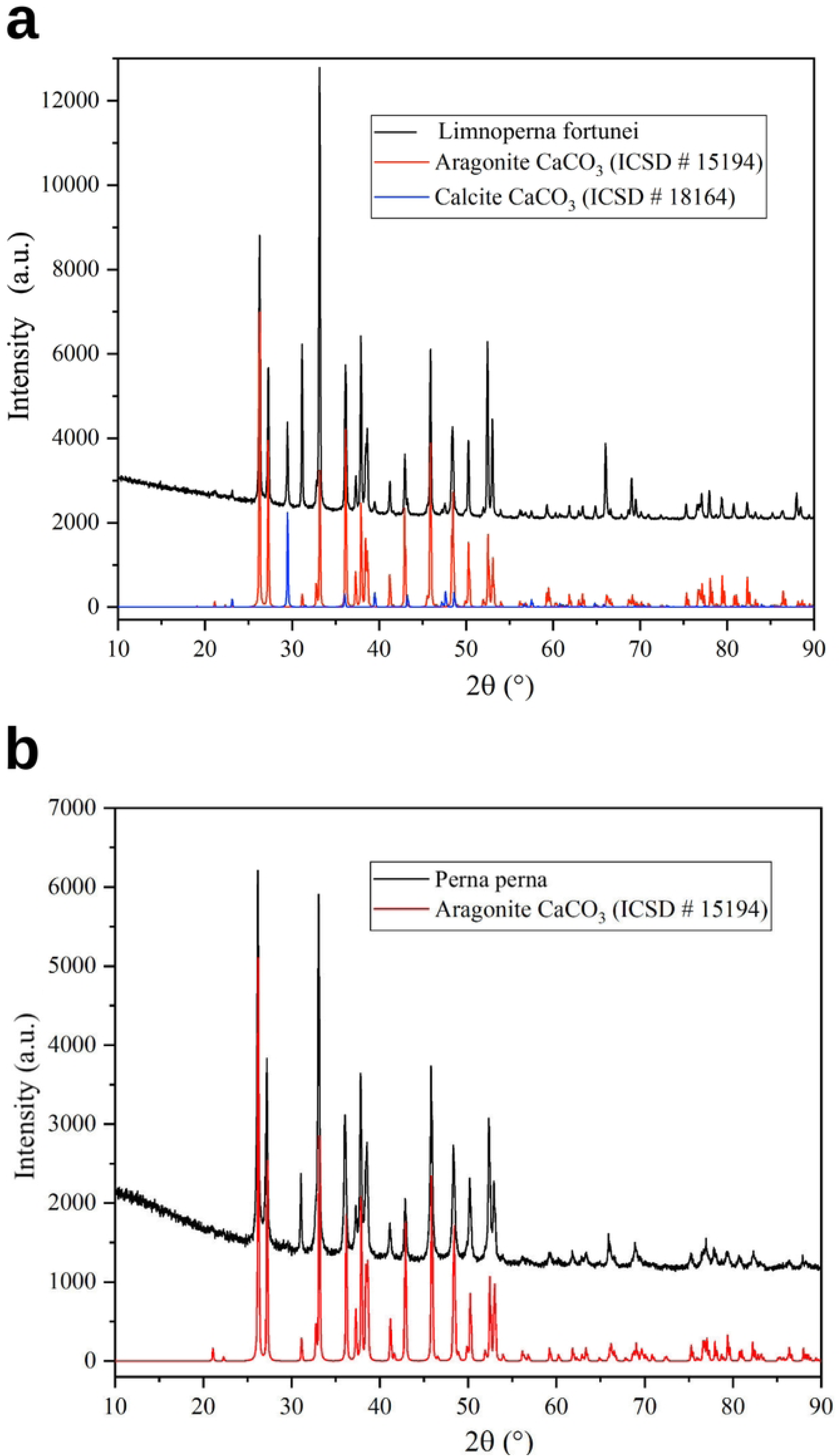
X-Ray diffractograms of mussels shells, (a) Aragonite and calcite phases are observed in the diffractogram of L fortune and (b) Aragonite is the only phase detected using XRD technique. No iron phase detected due to limits of detection.

**Table 2.** Phases, concentration (%), composition, crystalline system on the thermodynamic phases present on the shells of the freshwater bivalve L fortunei and the marine P perna mussels.

| Freshwater *Limnoperna fortunei* (Golden mussel) |  |  |  |  |
| --- | --- | --- | --- | --- |
| Phase | chemistry formula | Space group | Crystalline system | concentration (%) |
| Aragonite | CaCO<sub>3</sub> | Pmcn | Ortorrombic | 97.48 |
| Calcite | CaCO<sub>3</sub> | R3c | Trigonal | 2.52 |
| Marine *Perna perna* (Brown mussel) |  |  |  |  |
| Aragonite | CaCO<sub>3</sub> | Pmcn | Ortorrombic | >99.0 |

### Electron microscopy

Figures 5 to 9 show SEM and TEM images of magnetic powder obtained from demineralisation of the *Limnoperna fortunei* shell. The powder was attached to the magnetic stirring bar during the final step of demineralisation. The SEM data show the presence of calcium carbonate (CaCO_3_) along with iron (hydr)oxides nanoparticles, (Figure 5).

**Fig 5.**
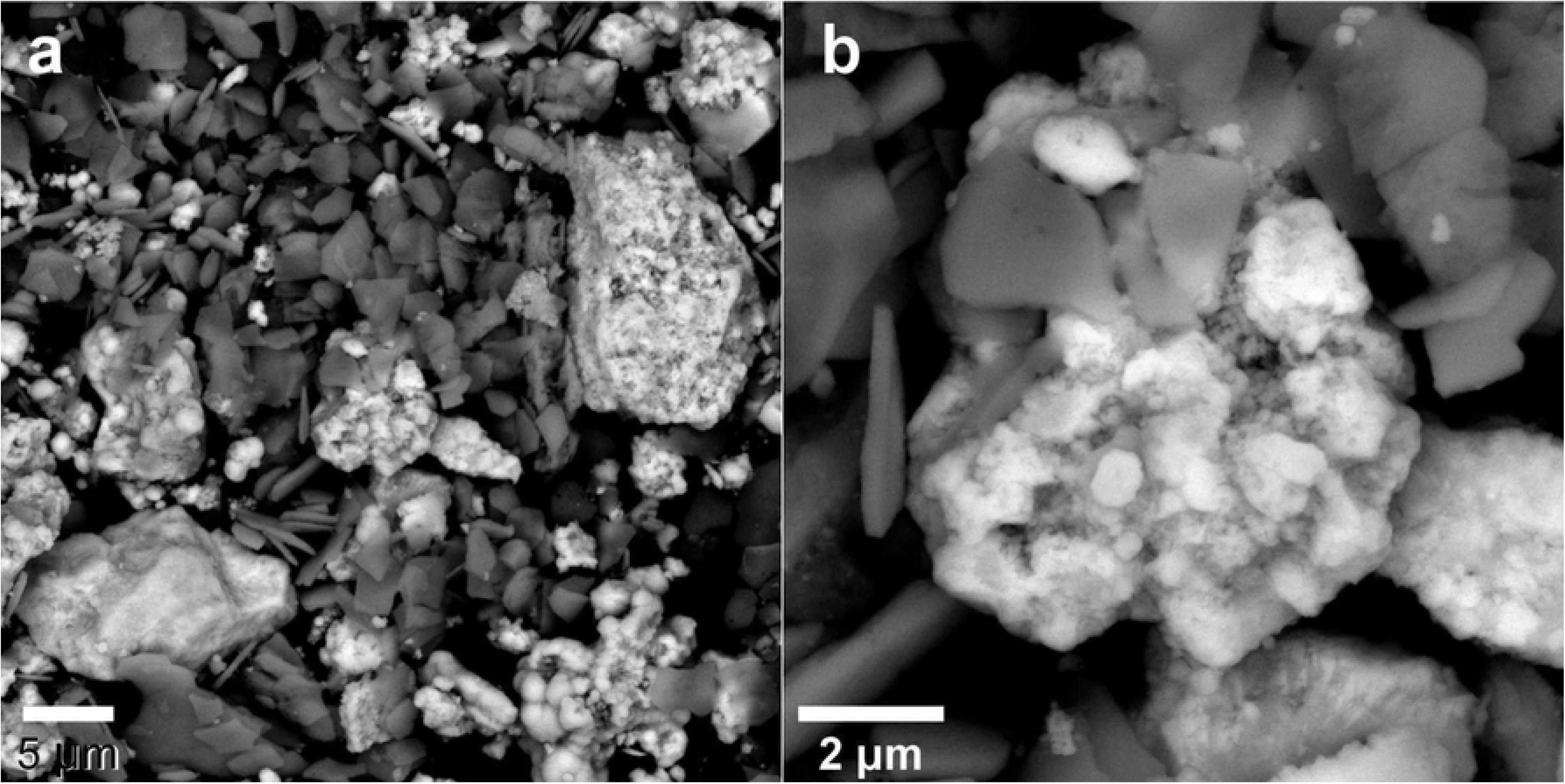
(a,b) Backscattering electron SEM images of the magnetic powder obtained from the demineralisation of the *Limnoperna fortunei* shell. The images show CaCO_3_ plates (grayish) along with iron (hydr)oxides (brighter particles). The iron (hydr)oxides appear as agglomerates of nanoparticles entangled with the CaCO_3_ plates as shown in image b.

These mixed phases were not separated by sonication as observed by TEM. Higher spatial resolution analysis of the magnetic powder was achieved by analytical TEM. The TEM data show either calcite or aragonite superimposed with the iron (hydr)oxides (magnetite, hematite, and goethite) (Figures 6 and 7).

**Fig 6.**
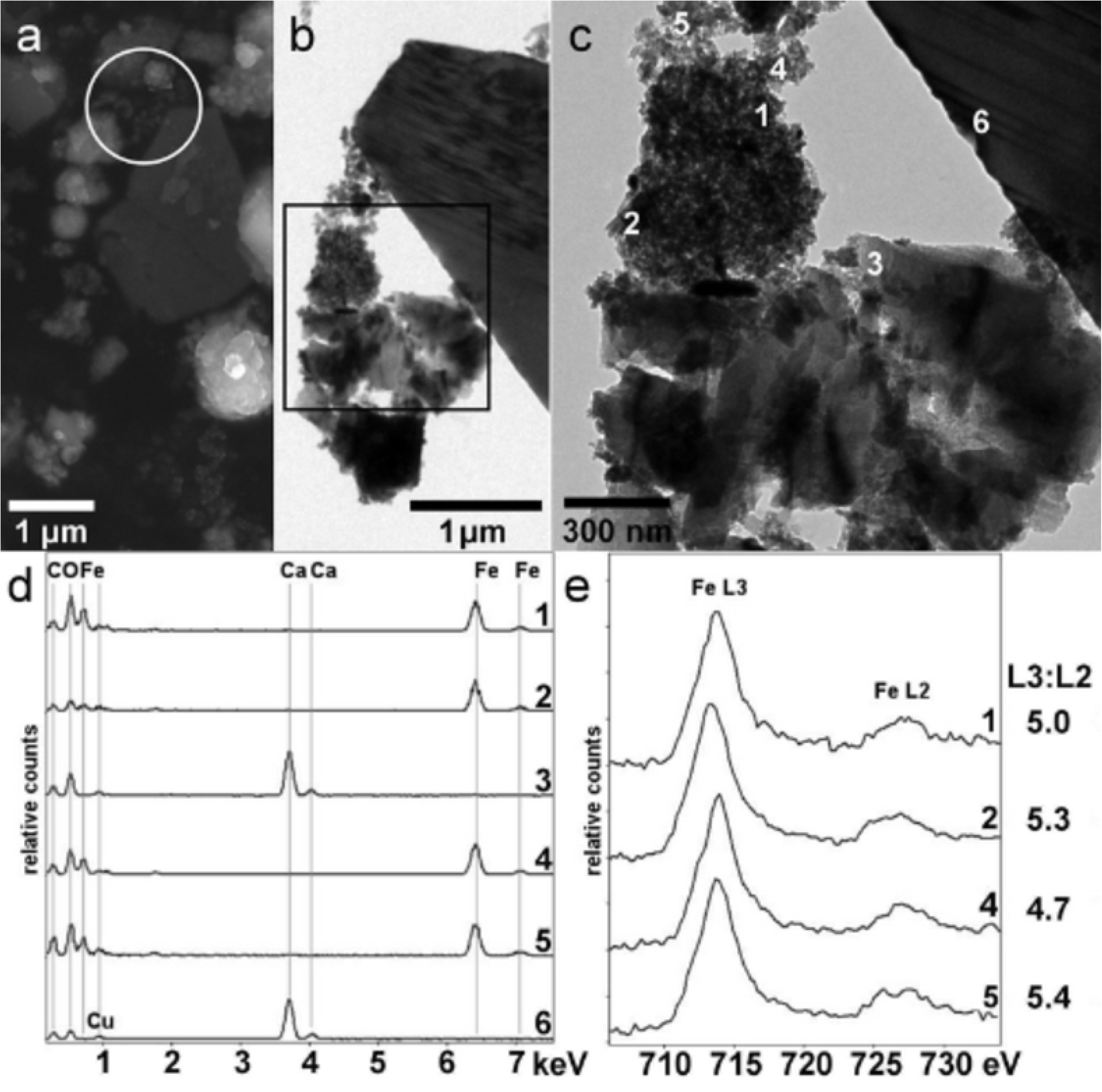
(a) Secondary electron SEM image and (b,c) bright-field TEM images of the magnetic powder obtained from the demineralisation of the *Limnoperna fortunei* shell. (d) EDX spectra and (e) EEL spectra at the Fe *L*-edges taken from the points marked in the image c. The *L*3:*L*2 values are the ratios between the intensities of the Fe-*L*3 and Fe-*L*2 edges, after background removal.

**Fig 7.**
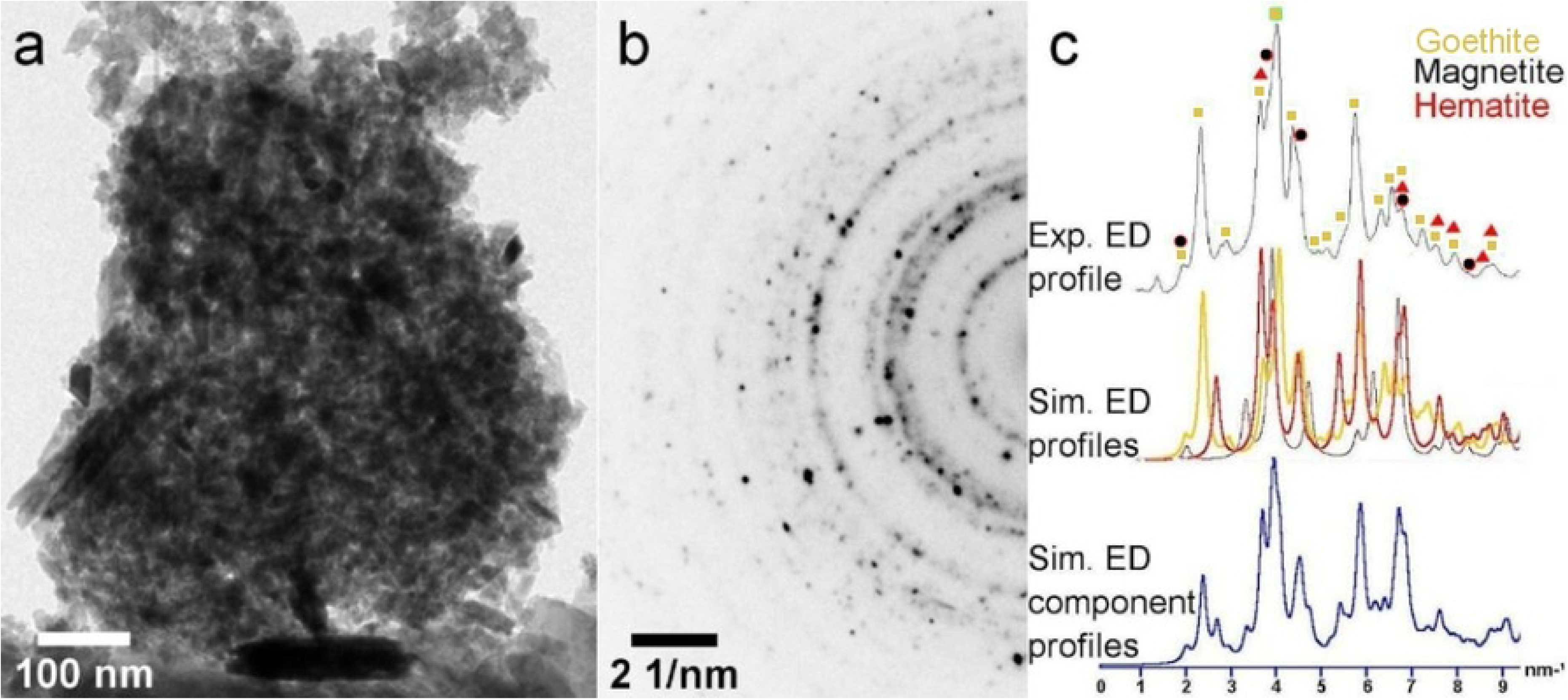
(a) Bright-field TEM image of an agglomerate of iron (hydr)oxide nanoparticles in the magnetic powder obtained from demineralisation of the *Limnoperna fortunei* shell. (b) SAD pattern (with an inverse contrast) of the agglomerate of particles shown in image a. (c) At the top, the experimental (Exp) electron diffraction (ED) profile of the SAD pattern shown in image b. At the middle, the simulated (Sim) ED profiles of goethite (green), magnetite (black), and hematite (red). At the bottom, the Sim. ED component profiles, considering the equal contents of goethite, magnetite, and hematite. The simulated ED profiles were performed with the help of the JEMS© software (v. 3.4922U2010).

These calcium carbonates and iron (hydr)oxide phases were confirmed by EELS/EDX (Figure 6.d-e) and SAD (Figure 7.b-c). In Figure 6.c the EDX spectra at points 4 and 7 indicate the presence of sole CaCO3. At points 1, 2, 4 and 5 (see Figure 6.c) EDX spectra indicate the occurrence of iron compounds. EELS analysis at the Fe *L*_3,2_-edges of spectra taken at these points were performed to confirm the presence of the magnetite. The white line ratios [Intensity (Fe-*L*_3_): Intensity (Fe-*L*_2_)] at points 1 (5.0), 2 (5.3) and 4 (5.4) agree with the data reported for magnetite (5.2±0.3) in the literature [24]. Figure 8 shows the TEM image, SAD pattern and EEL spectrum of a sole magnetite nanoparticle found in the magnetic powder. Again, the magnetite and other iron oxide phases are mixed with either aragonite or calcite in the magnetic powder sample. We attempted to localize the magnetic nanoparticles in a cross-section of the *Limnoperna fortunei* shell, but it was unsuccessful. We have not found any iron oxide particle at the periostracum layer nor in the calcite crystals just after the periostracum.

**Fig 8.**
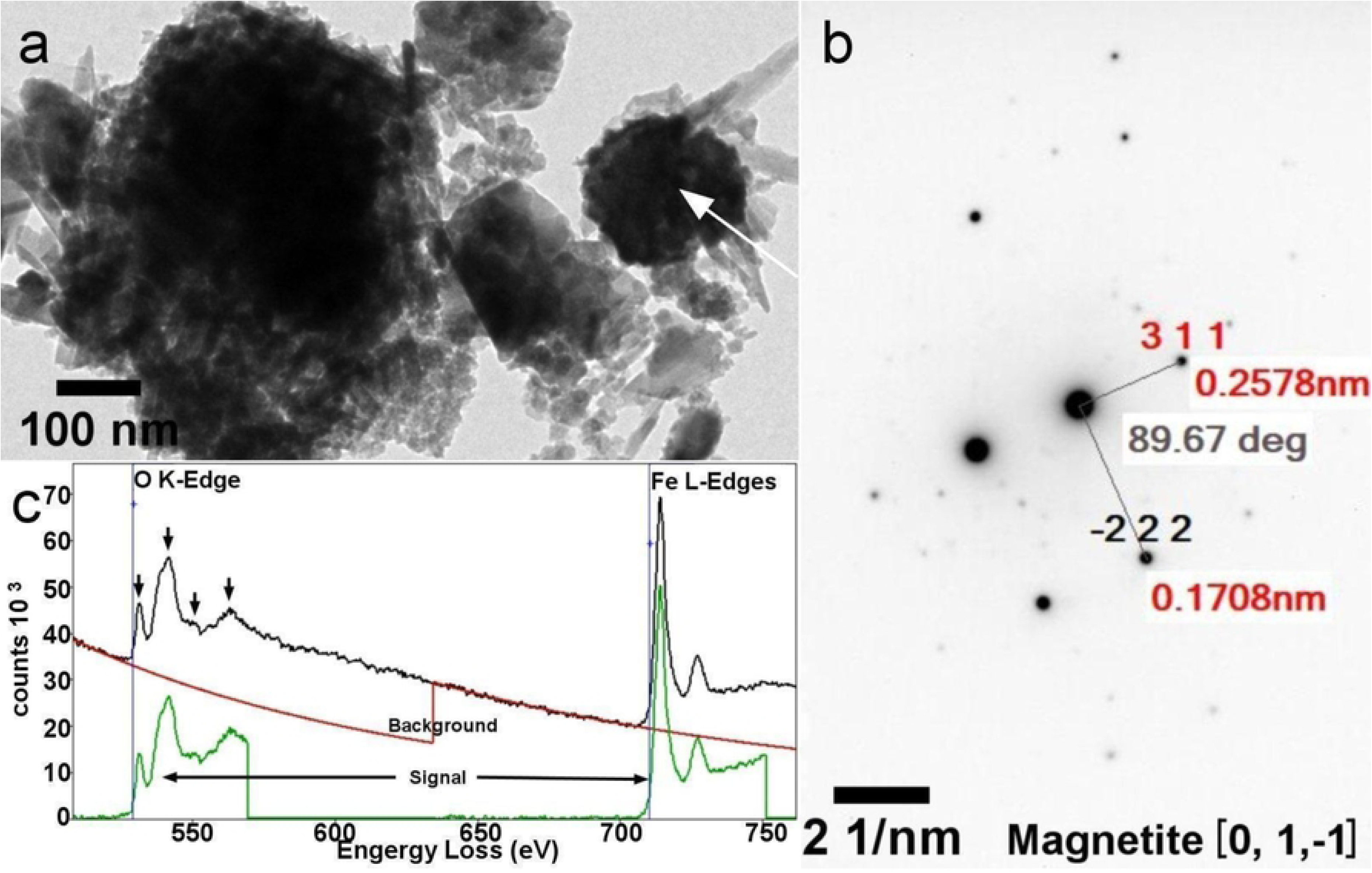
(a) Bright-field TEM image of an agglomerate of iron (hydr)oxide nanoparticles in the magnetic powder obtained from the demineralisation of the *Limnoperna fortunei* shell. (b) SAD pattern (with an inverse contrast) of the magnetite particle pointed with the arrow in image a. (c) EEL spectrum at the O *K*-edge and Fe *L*-edges of the magnetite nanoparticle pointed in image a. All the four characteristic peaks at the oxygen *K-*edge for magnetite are indicated with the black arrows.

Nevertheless, roughly round nanoparticle aggregates of iron (hydr)oxide (likely goethite) were found in the nacar layer (Figure 9), associated with aragonite. Figure 9 shows the SEM and TEM images of a portion of the nacar layer with an iron-rich nanoparticle attached to it. TEM and especially HRTEM (Figure 9.c) indicated that the 50-100 nm round particles are in fact aggregates of 5-10 nm nanoparticles. Fast Fourier Transformed (FFT) analysis of the HRTEM image confirms that the 5-10nm nanoparticles are goethite.

**Fig 9.**
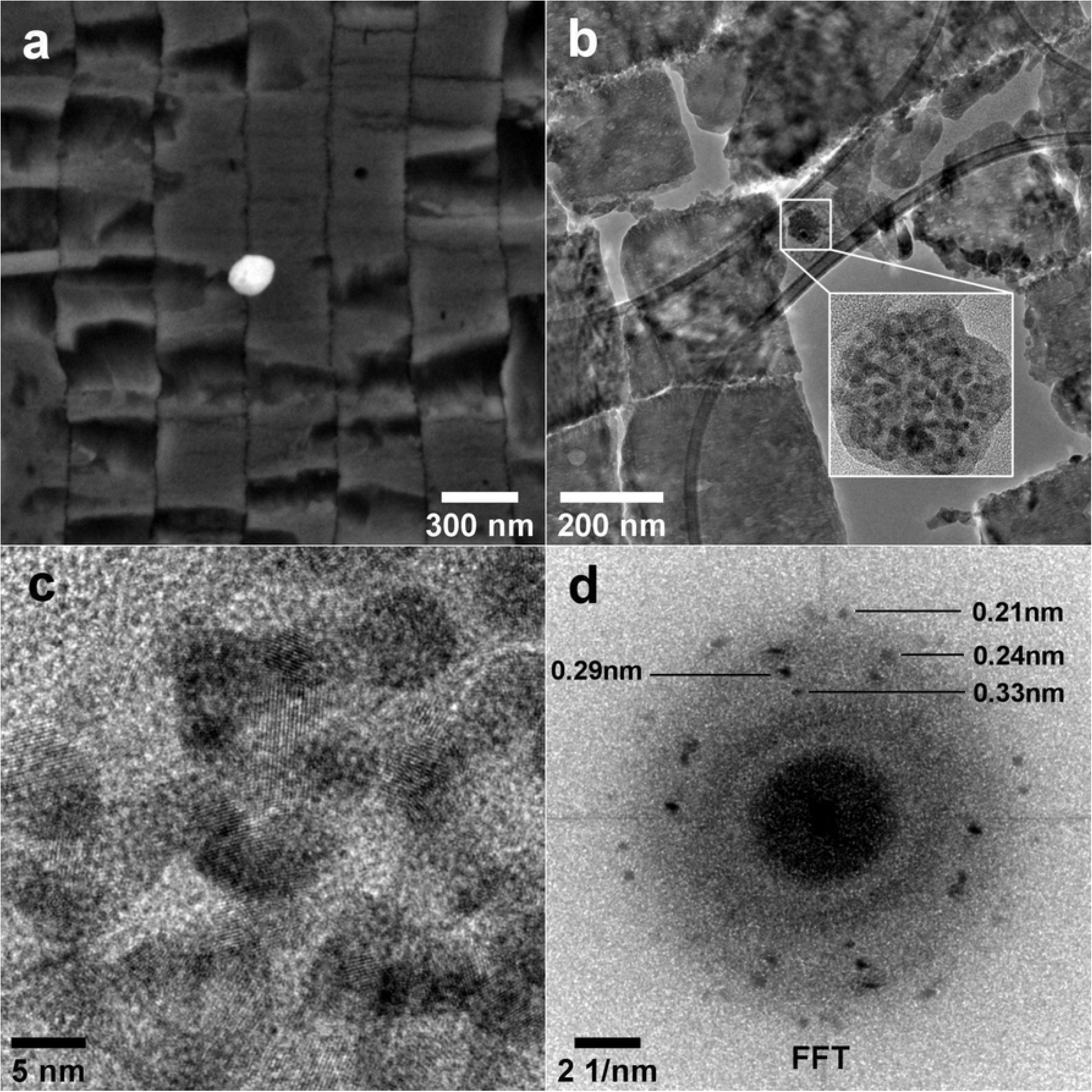
SEM and TEM images of the cross-section of the nacar layer of the *Limnoperna fortunei* shell. (a) Backscattering electron SEM image showing a round iron-rich nanoparticle attached to the aragonite tiles. (b) Bright-field TEM image of a thin cross-section layer (100 nm thick) showing a round nanoparticle aggregate in between the aragonite tiles. (c) HRTEM image showing in detail the crystalline nanoparticles of the aggregate marked in b. (d) Fast Fourier Transform of the HRTEM image. The distance of the lattice fringes is indicated and agrees with goethite.

### Raman Spectroscopy

The magnetic powder collected from demineralized shells was analyzed by Raman spectroscopy. Some of the Raman spectra are shown in Figure 10. Magnetite, goethite, and other iron-oxy(hydr)oxides were found. Spectrum 1 was acquired at a dark point seen in the light optical image (not shown). It shows the line 666 cm^-1^ characteristic of magnetite. In a yellowish region of the sample seen at the microscope we found goethite (Figure 10 - spectrum 2). Mixed phases were also observed at some points as can be seen in the spectrum 3, which shows the presence of both magnetite and goethite. Aragonite (Figure 10 - spectrum 4) was found in the magnetic powder samples. These results corroborate the findings in the SEM and TEM analyses. During the Raman measurements, we observed the formation of the hematite under the laser exposure at some points, so the laser intensity was adjusted to avoid it.

**Fig 10.**
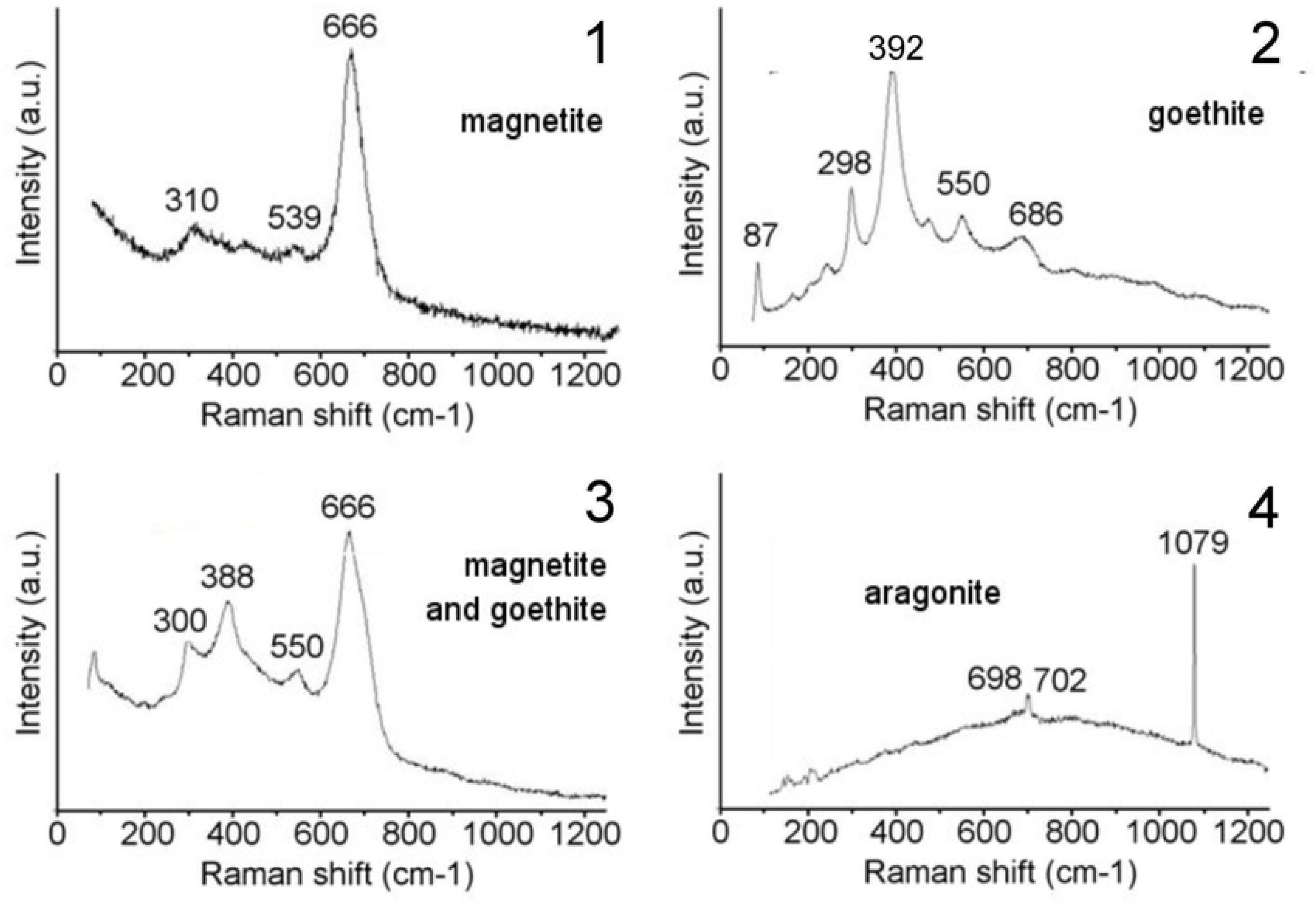
Raman spectra (1,2,3 and 4) of the magnetic powder obtained from the demineralized Limnoperna *fortunei* shell.

Additionally, Raman spectroscopy was employed to confirm the occurrence of magnetite particles, collected from just grounded mussel shells of P perna (11.a) and L fortunei (11.b) using porcelain mortar and pestle.

**Fig 11.**
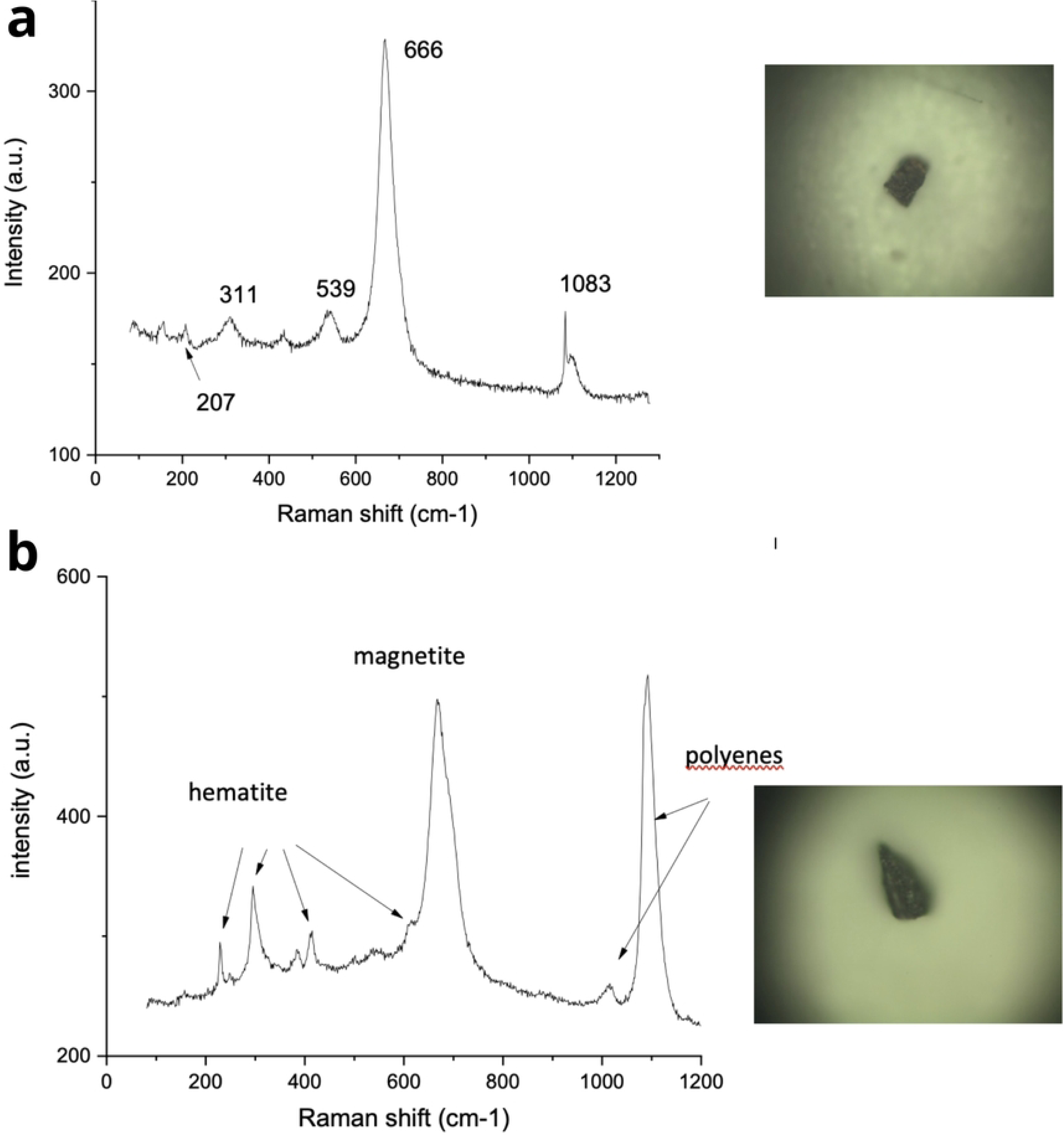
(a) Raman spectra of the magnetic particles obtained from the Perna perna shell. The spectrum shows magnetite (Fe_3_O_4_) lines at 666, 540, 311 cm^-1^, and aragonite (CaCO_3_) lines at 1083, 207 and 155 cm^-1^. Particle size ∼ 100 x 150 um (image above right, 10X Objective); (b) Raman spectra of the magnetic particles from L fortunei shell. The spectrum shows magnetite, hematite, and polyenes peaks. Particle size∼ 25 x 50 um (image above right, 50X Objective). Powder obtained from fresh crushed (in a porcelain pestle and mortar).

Also using Raman to analyze sections of gills of freshwater L fortunei we identified a variety of micrometer size particles of hematite, lepidocrocite, aragonite, calcite and anatase (Figure 12). In other soft parts of the body of freshwater *L fortunei* we observed the presence of microparticles of hematite. There is the possibility that these microparticles captured by the gills might arrive inside the circulatory system of the mussel as we have observed on many occasions using Raman.

**Fig 12.**
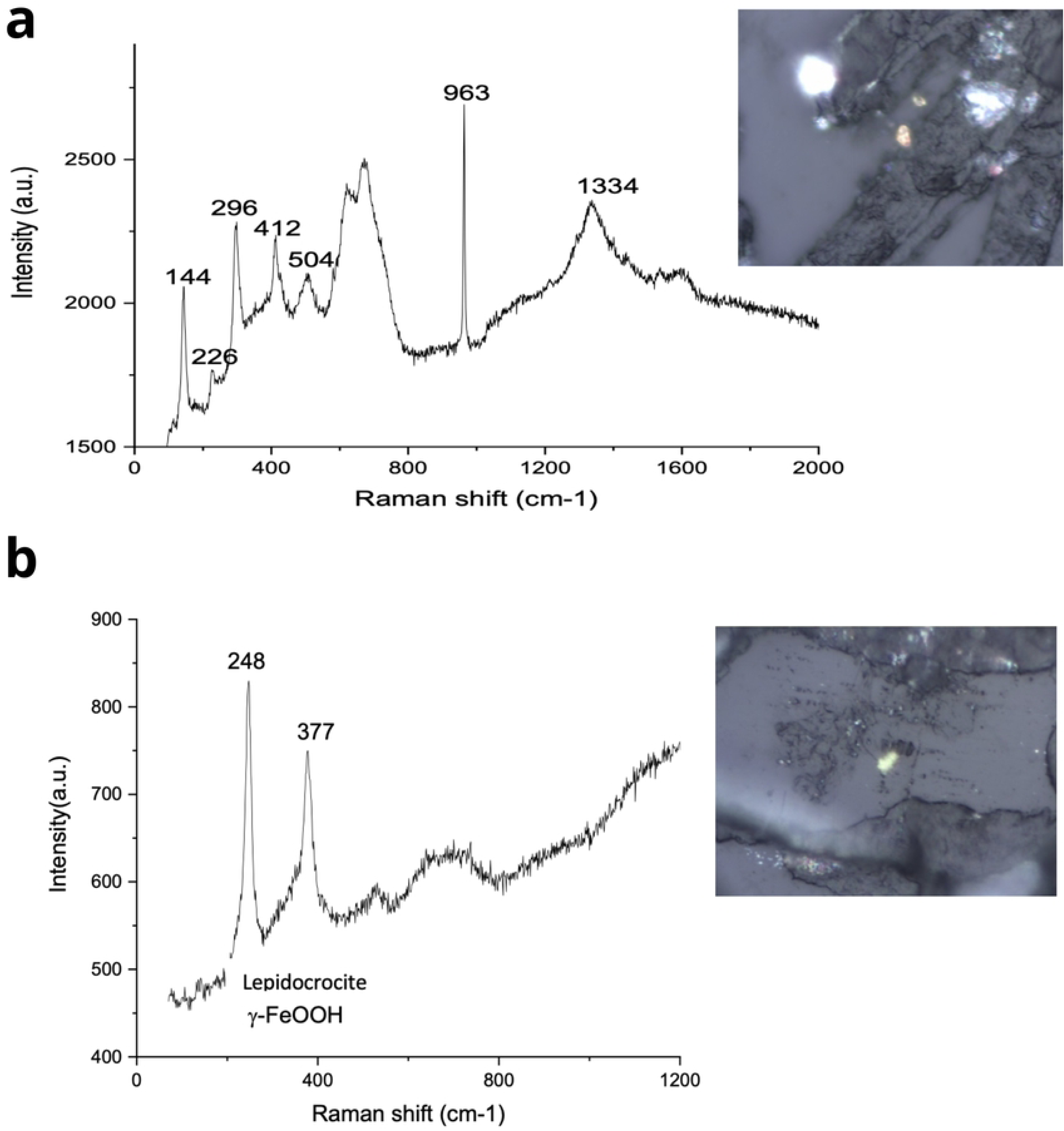
(a) Raman spectra of a section of the L fortunei gills [30]. Spectrum above have line 144cm^-1^ (anatase TiO_2_); lines 226, 296, 412, 610 cm^-1^ (α-Fe_2_O_3_, hematite). The 963cm-1 line may be of apatite, a calcium phosphate mineral; (b) Spectrum above have lines 248cm-1, 377 for (gamma-FeOOH, lepidocrocite). Laser at 632.8nm, CCD cooled to -126 °C, 100X lens objective.

### Vibrating-Sample Magnetometer (VSM)

To measure the magnetic properties of the shell of the Perna perna and Limnoperna fortunei we performed magnetometry Vibrating Sample Magnetometer (VSM).

In the P *perna* shell fragments we can observe the predominant signal of diamagnetic materials from amorphous calcium carbonates as observed by SEM. Although the magnetic signal observed in Figures 13.s1, 13.s6, 13.s7, 13.s8 and 13.s9 are relatively low, we can still notice small magnetic hysteresis referring to the presence of a magnetic material in the mussels’s shells, for example, magnetite (Fe_3_O_4_) has ferrimagnetic order. The other fragments in Figures 13.s2, 13.s3, 13.s4 and 13.s5, on the other hand, have a practically diamagnetic signal.

**Fig 13.**
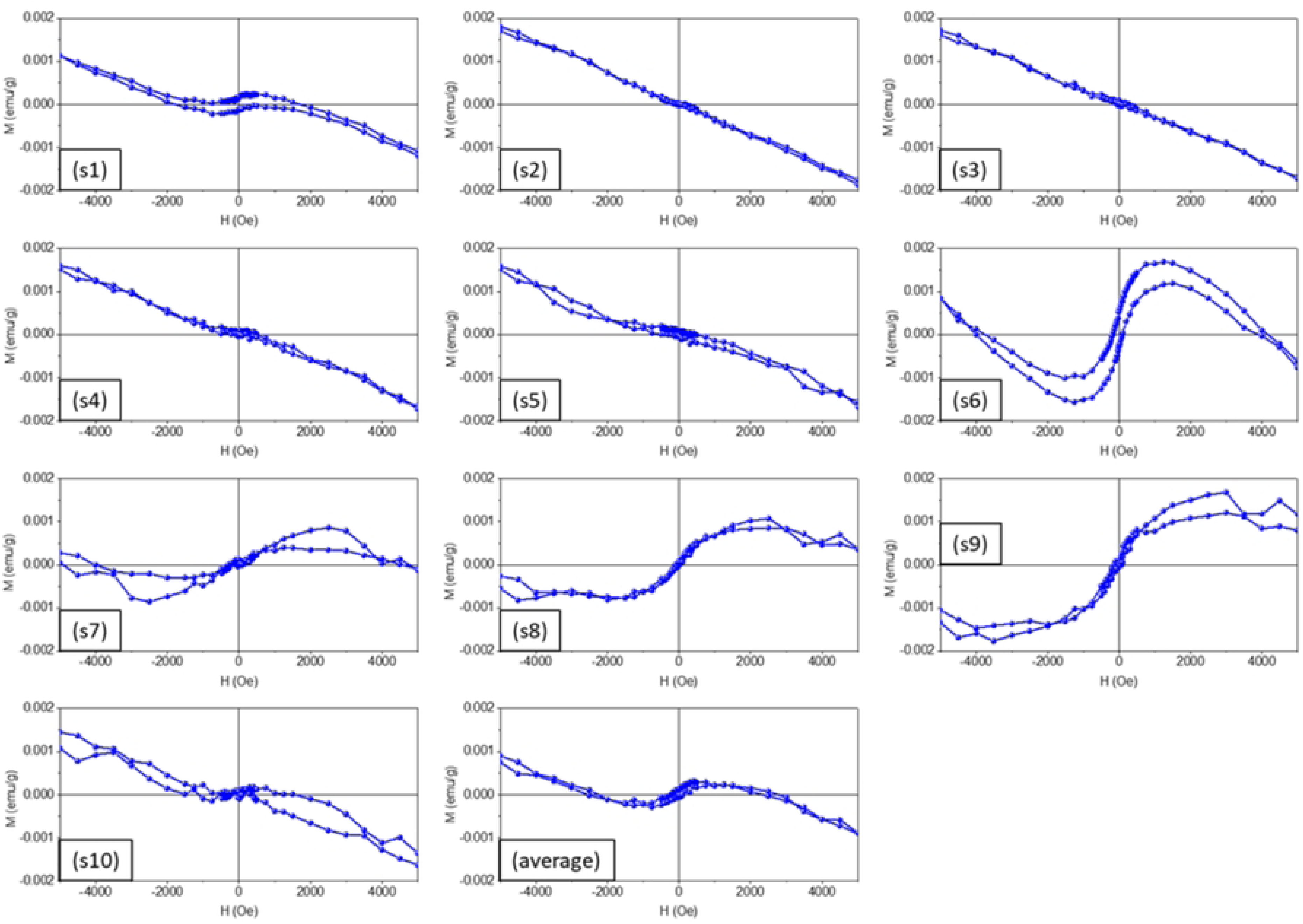
Magnetization M (emu/g) versus magnetic field strength H (Oe) measured in 10 (ten) pieces of the shell of marine mussel Perna *perna* (Brown mussel). Pieces were numbered s1 to s10 and the average of assays is presented in the last graph (bottom, middle).

In the magnetic curves that present the highest concentration of magnetite, we still have the curves with low coercivity and magnetic remanence, highlighting the fact that these magnetic materials present in the shells are in low concentration and present a superparamagnetic behavior characteristic of materials of nanometric sizes.

However, as magnetite (ferrimagnetic), hematite (antiferromagnetic) and/or goethite (antiferromagnetic) nanoparticles can be present in these fragments and competing with each other; and therefore prevailing the one with the highest intensity due to the volumetric measurement of the magnetometer VSM [25].

For the L *fortunei* shell VSM test, we divided the shell into fragments and numbered them 1 to 10 as shown in Figure 14, we can observe again that the shells have a ferrimagnetic character (FI) for the fragments of Figures 14.s1, 14.s6 and 14.S9, in addition to other components: diamagnetic (DM) Figure 14.s2, 14.s4 and 14.s5 and paramagnetic (PM) present in Figures 14.s3 and 14.s7.

**Fig 14.**
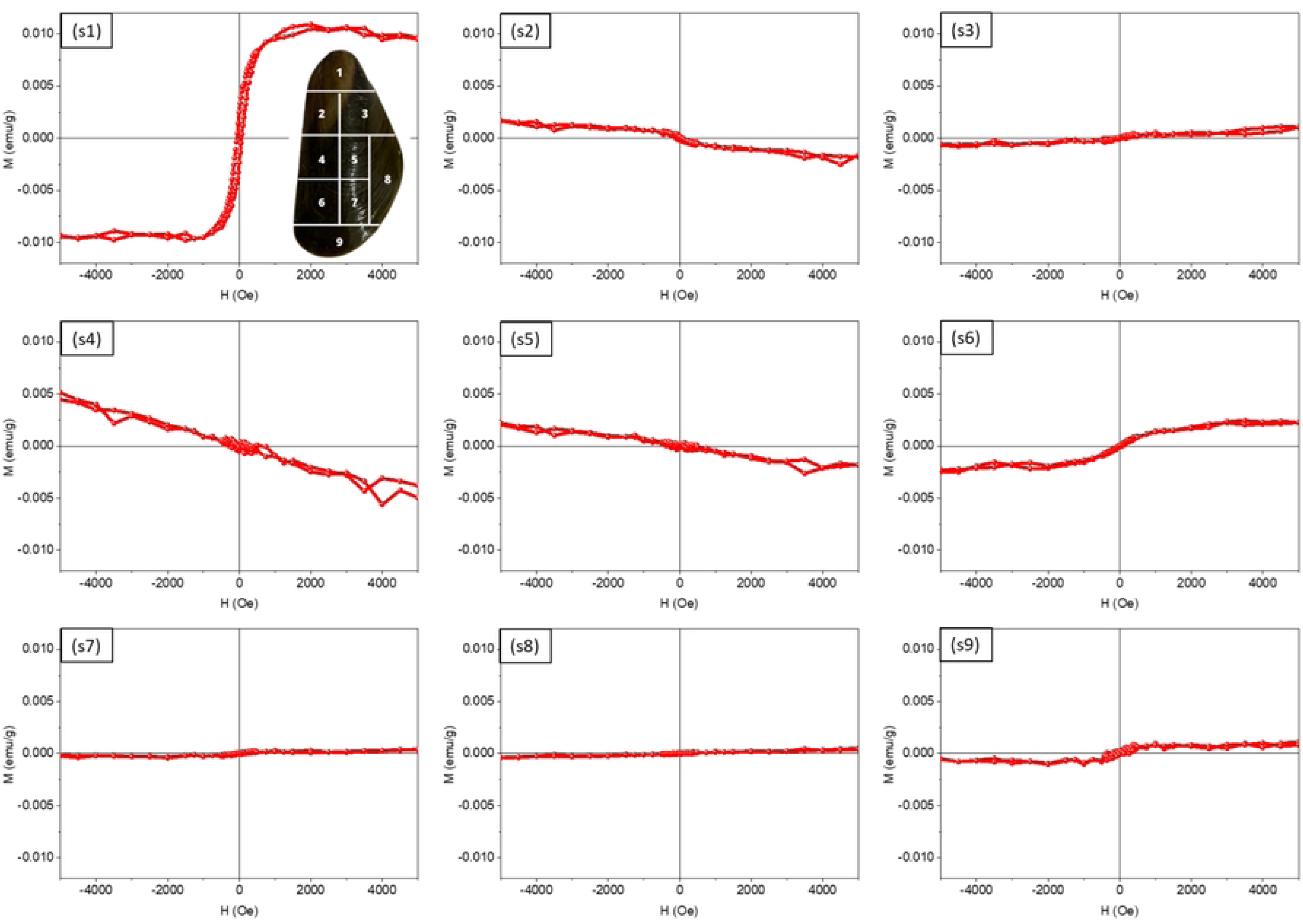
Magnetization M (emu/g) versus magnetic field strength H (Oe) measured in 9 (nine) pieces of the shell of freshwater mussel L *fortunei (*Golden mussel). Pieces were numbered s1 to s9 (see image inside run s1, top right).

It is worth noting that the samples have low magnetization due to low iron concentration, and thus PM, DM and FI signals overlap in this concentration range. Pure iron material has about 221 emu/g [26], while magnetite nanoparticles obtained through chemical/physical processes have an average value of 80 emu/g [27, 28]. The saturation magnetization, from lower to higher signal (FI) can be estimated to have a concentration of 10 to 150 ppm of magnetite in the mass of these fragments analysed.

Considering that the sensitivity of the VSM-7400 equipment is around 1x10^-7^ emu and that the technique is volumetric, accounting for all magnetic signals present in the samples or their absence, some of these fragments may have magnetic nanoparticles but the total signal is masked by the signal from other non-magnetic components.

Figure 15 shows the saturation magnetization (Ms) of collected magnetic samples from a simply sanding of L *fortunei* shells. The Ms value 0.21 emu/g is small but measurable as expected due to the small mass of magnetic particles sanded from the mussel shell.

**Fig 15.**
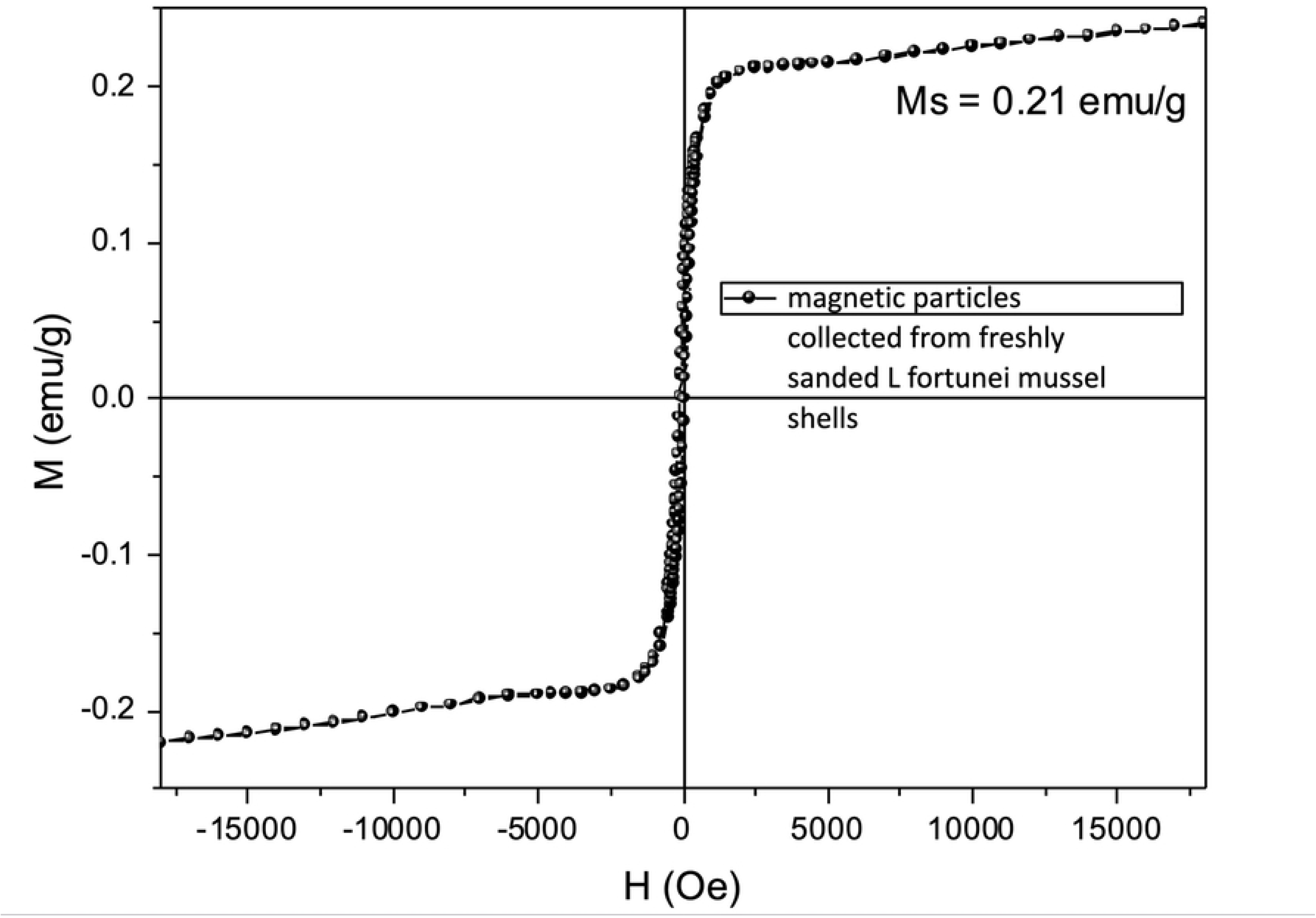
Magnetization M (emu/g) versus magnetic field strength H (Oe) measured in magnetic particles collected from simply sanded shells of freshwater mussel Limnoperna *fortunei (*Golden mussel). Commercial sandpaper (SiC) and NdFeB magnet were used in the experiment. See Supporting S1- Video: Sanded L fortunei mussel shell: collecting magnetic particles. Cleaned and dried Limnoperna fortunei shell were sanded with commercial sandpaper (SiC).

## Discussion

Are the nanomagnetic particles present on the shell of *L fortunei* and *P perna* just ingested particles or absorbed by the gills?

Mussels cycle liters of water through their bodies and magnetic nanoparticles could come from microalgae feeding [30] or present in the water being continuously filtered. Then they could diffuse into hemolymph, ending up eventually in the shell itself. Magnetic iron-oxides are found in a multitude of environments worldwide simply by natural occurrence as well as by human made environmental pollution.

Bivalves continuously filter microparticles and nanoparticles of many different compositions and minerals both at the river and at the sea, the most abundant being silica and aluminosilicate clays. But none of those appear in noticeable amounts in the shell composition as the ICP-OES (Table 1) show. Previous studies [31, 32] have arrived at the same conclusion, that the presence of nano or microparticles of different compositions does not mean that they will be incorporated in the shell. For this reason, the composition of the shell is quite exclusively made of CaCO_3_, chitin and proteins.

Raman spectroscopy (Figures 11 and 12) indicates that many microparticles were detected in the body of L fortunei, even magnetite microparticles. But the amount of iron detected by ICP-OES on the shell is much lower than 1%, like other heavy metals detected.

This is quite understandable considering that any particle could only arrive at the extrapallial surface after traversing the Mantle. The Mantle is a rather complex gelified structure [33], that somehow selects, via "filtering", dissolution and precipitation, the inorganic particles that will arrive at the extrapallial external surface. Therefore many microparticles and surely many more nanoparticles diffuse and are present in the soft body of the bivalves, which is new information, but these are not freely able to cross and arrive at the outer face of the Mantle. The Mantle is not permeable, quite the contrary. MEV-EDS images of the mantle of the L fortunei [34] show a quite complex structure, fully of calcium associated with phosphate.

Since Aristotle [35] the design and the growth of the mollusk shell has attracted the attention of savants and researchers across many fields and for many centuries. But so far there is no detailed description that has been universally accepted. Curiously the metal and metal compounds content of mollusk shells are still scarcely studied, albeit the fact that it could provide decisive and useful information in the understanding of the biomineralization process.

The interference of iron cations on the crystallization, nucleation and growth of CaCO_3_ thermodynamic phases are presented in incountable scientific works in many areas. One line of explanation is based on the cation radius [31, 32]. Calcium has a Pauling ionic radius of 99pm and in calcite, an ionic crystal, the Ca atoms have a face-centered distribution and the structure, viewed as a rhombohedron, have the triangular CO_3_ groups lying at the centers of each edge of the rhombohedron. This results in layers of CO_3_ groups lying normal to the c-axis of calcite, with layers of Ca atoms lying between them. Iron ions (Fe^+2^ and Fe^+3^) with radii less than calcium ion would replace calcium as iron carbonate (siderite) is isostructural with calcite. According to this line of argument not only iron but many divalent metal cations smaller than calcium could substitute Ca^+2^ [36].

Studies on the interference of iron on the crystallization of CaCO3 has immediate economic importance as the scales of CaCO_3_ occludes pipes in water treatment systems, desalination plants, mineral, and crude oil pipelines, etc.

Contrary to the above suggestion (easy replacement of Ca by Fe ions) many experimental studies [37] agreed that, in an aqueous phase, and in the presence of minute amounts of ferrous iron Fe^+2^ calcite will not occur at all or only marginally. Calcite phase is the most stable form of crystalline calcium carbonate and very difficult to dissolve as soon as it nucleates and grows at the walls of pipelines. However ferrous ion presence does not prevent the nucleation and growth of aragonite. The reason for this behavior is not yet clearly understood but it has been observed since the problem caught the attention of scientists [37, 38].

Molecular Dynamics studies [39] also advocate that calcite growth is inhibited in the presence of ferrous ions and other divalent metal ions smaller than calcium. Initially ferrous ions incorporation at the step growth of calcite is energetically favorable (exothermic) but as the incorporation continues it produces mismatches that ultimately turns the growth of the crystal energetically unfavorable (endothermic) and stops it.

On the other hand, ferric ion Fe^+3^ too influences CaCO_3_ nucleation and growth. As Fe^+2^ is oxidized to Fe^+3^, even in very small amounts, the calcium carbonate present tends to crystallize in the calcite phase as many works have noticed. One suggestion for this to happen is because iron electronegativity (1.8) is much higher than calcium (1), so if iron and calcium coexist the crystallization of calcium is directly affected. Evidently other variables might interfere, especially acidity or basicity.

Recently it has been reported that the acidification of sea waters not only tends to corrode bivalve shells but also alters shell structure with the aragonite nacre layer thickening at the middle of the shell while being wiped out from the acidic borders [40]. Aragonite is more easily dissolved, especially in acidic waters. But at the center of the shell, it is not in contact with acidic waters and the thickening increases the mechanical strength of the shell. Therefore, it might be reasonable to think that the magnetic particles we have encountered in both maritime and freshwater bivalves are related with the aragonitic nacre layer. How? it is a matter of further studies.

Finally, the interference of Fe ion on the crystallization of CaCO_3_ is based on the reasoning that the concentration of Fe should be in the ppm (mg/L) range which was confirmed by ICP assays presented here.

One hypothesis discussed by A Mount et al [41] is that calcite nucleates and grows inside hemocytes, at least in oysters. For the iron nano-compounds (oxide or hydroxide) to work as nucleating agents this could be more easy and less costly (in energy terms) for the hemocytes if nucleation occurs inside the cell compartment. Another possibility is that granular hemocytes delivering ACC and iron nanoparticles aggregate at the extrapallial surface. The pH will define the nucleation phase as Fe^+2^ will favor aragonite nucleation and Fe^+3^ close to the external environment will favor calcite formation. Both possibilities help to understand how the occurrence of CaCO_3_ crystalline probably occurs by combining organic matrix induction and the nucleating agent oxidation state, as it happens on scale formation.

The list of animals with a proven presence of magnetic particles continues to rise. However, to date the presence of magnetic nanoparticles *in the shell of molluscs* has not been reported. Scientific articles published so far focus on the most diverse uses for the CaCO_3_ present in shells [5]. However, to obtain the shell powder a heat treatment of up to 1000 °C or higher [6] is performed, oxidizing the nanoparticles into Fe_2_O_3_. Also, the presence of magnetite appears to occur in human organs including the human brain [42, 43]. More recently, an external source of magnetite nanoparticles present in the human brain was presented by researchers as being the result of urban pollution [44]. According to the study, what differentiates the two types would be the morphology.

Our own findings are evidence of the magnetic particles present at the freshwater bivalve *Limnoperna fortunei* and saltwater *Perna perna* shells. The TEM results show magnetite and maghemite along with other Fe-(hydr)oxides nanoparticle aggregates close to the microsize plates of aragonite. Raman spectroscopy results showed the occurrence of magnetite, goethite, other iron compounds and minerals microparticles, along with aragonite particles. Results shown here helps to understand why the occurrence of magnetite particles have not been noticed so far. Being nanoparticles, they easily oxidize to other iron oxides.

Considering a 10 nm sphere iron-rich particle and considering a 300 miligrams *Limnoperna fortunei* shell weight, we arrive at an astounding number of iron-rich nanoparticles, around 2 x 10^17^ particles. If they were all magnetite or maghemite, they could only be a part due to the magnetic tendency to aggregate (See Figure 2). Therefore, it is still a matter of importance to clarify if magnetite biomineralization is concurrent with ACC as already reported for some bacteria [45]. Could the biogenic formation of ACC somehow help in the nucleation and growth of magnetite or is it the other way round?

The Ca^+2^ and Fe^+2^ ions are toxic to the cells and the nucleation and growth of calcium carbonate and iron oxide are a matter of survival for the cells. But many studies have suggested that ACC nanoparticles with few nanometers in size are available in the water [46, 47] and probably captured by the gills. Results from Raman spectroscopy presented here does show a variety of mineral microparticles inside the soft body and this could be the case for the magnetite nanoparticles too as the phase change from iron (oxy)hydroxides (ferrihydrite) to iron oxides have been reported for the chiton teeth.

### Iron inside the cells

Most magnetic materials (paramagnetic and ferromagnetic) in organisms are composed of iron, especially Fe oxides. In practice all organisms require Fe to function normally. This is mainly due to the reducing activity of Fe which allows it to play an important role in energy processes. In organisms, iron is stored in the ferrihydrite phase (5Fe_2_O_3_.9H_2_O), this phase bound to the ferritin protein.

Heinz A. Lowenstam [1], in 1962, was the first to report the presence of biochemically precipitated magnetite (biogenic) in the covering of the teeth of the radula (lingual plate) of the chiton (marine molluscs of the Polyplacophora class). Prior to this discovery, magnetite was thought to form only in igneous or metamorphic rocks under high temperatures and pressure. According to recent studies this iron is deposited in the ferrihydrite phase and nucleated in the protein mesh, forming one or two distinct rows of reddish teeth. Subsequently, ferrihydrite is rapidly transformed into magnetite through an unknown process.

So far there is experimental evidence of the presence of magnetite nanoparticles-MNP in bacteria, chitons, fish and humans [48]. The so-called “Magnetoreception”, that is, spatial orientation by terrestrial magnetism, is a subject of scientific investigation [49]. Understanding the origin of the formation of magnetite nanoparticles is still a matter of discussion in the scientific field and some experts on the subject suggest that the formation of magnetite nanoparticles may have occurred before the emergence of eukaryotic cells [50].

The results obtained here indicated that magnetic particles occurs in very small particles, nanometers in size, these biomineralized magnetic nanoparticles have approximately equal volume and shapes and might be considered as a source of nanoparticles for Medicine and Molecular Biology [51,52,53].

Recent advances in the comprehension of role of hemocytes in the biomineralization of the shell [54, 55] has indicated that part or the totality of calcium that arrives at the mantle arrives as calcium carbonate (CaCO_3_) and possibly [56] in the amorphous state (ACC). According to these findings the CaCO_3_ nucleates and grows inside a fraction of the motile hemocyte cells and then are transported to the mantle.

The energetic status of these aggregates is very important to be considered. Thermodynamically ACC have an excess of energy and are metastable. These sub-micrometer size aggregates are then delivered to the extrapalial space where they are added to the inner faces of the growing shell. There, as it is suggested, it changes to crystalline phases (calcite and aragonite). This transformation of amorphous-crystalline is exothermic and tends to volume contraction as crystalline occupies less volume than the amorphous phase.

The concomitant biomineralization of magnetite and calcium carbonate have been recently reported in magnetotactic bacterias [45] and the authors believe that biomineralization of more than one mineral phase could be widespread [57]. While the carbonate is amorphous (ACC) the magnetite is crystalline. While the ACC particles are micrometers in size the magnetite are nanometer sized. If this is the case, magnetite nanoparticles are distributed throughout all shell layers.

## Conclusions

The dynamics of biomineralization, that is the steps and sequences of events on the construction of the shell of bivalves, rest today unresolved. The more established theory advocates for the main role of proteins and polysaccharides (chitosan) that produce a composite structure together with proteins, that are the nucleating sites for the inorganic substances to nucleate and grow. More recent advances [58, 59] suggest that carbon and phosphorus compounds and water have a prominent role as they self organize and control the growth of the shell. From this perspective the presence of proteins and polysaccharides are the intervention of the genome to improve and increase the strength of the shell as they create fibers and glues to construct the biogenic composite.

In this work we have added a new element, the magnetic iron compound magnetite (FeO.Fe_2_O_3_) and/or maghemite (gamma-Fe_2_O_3_), that could play a role in the process of self-organization and construction of the shell. To be sure that magnetite/maghemite on the shell was not simply a singularity of the freshwater exotic species bivalve Limnoperna fortunei we identified magnetic particles on the marine bivalve Perna perna shell.

To be sure that the magnetic particles were not an artifact of the demineralization process we performed the sequence present in the diagram of Figure 16.

**Fig 16.**
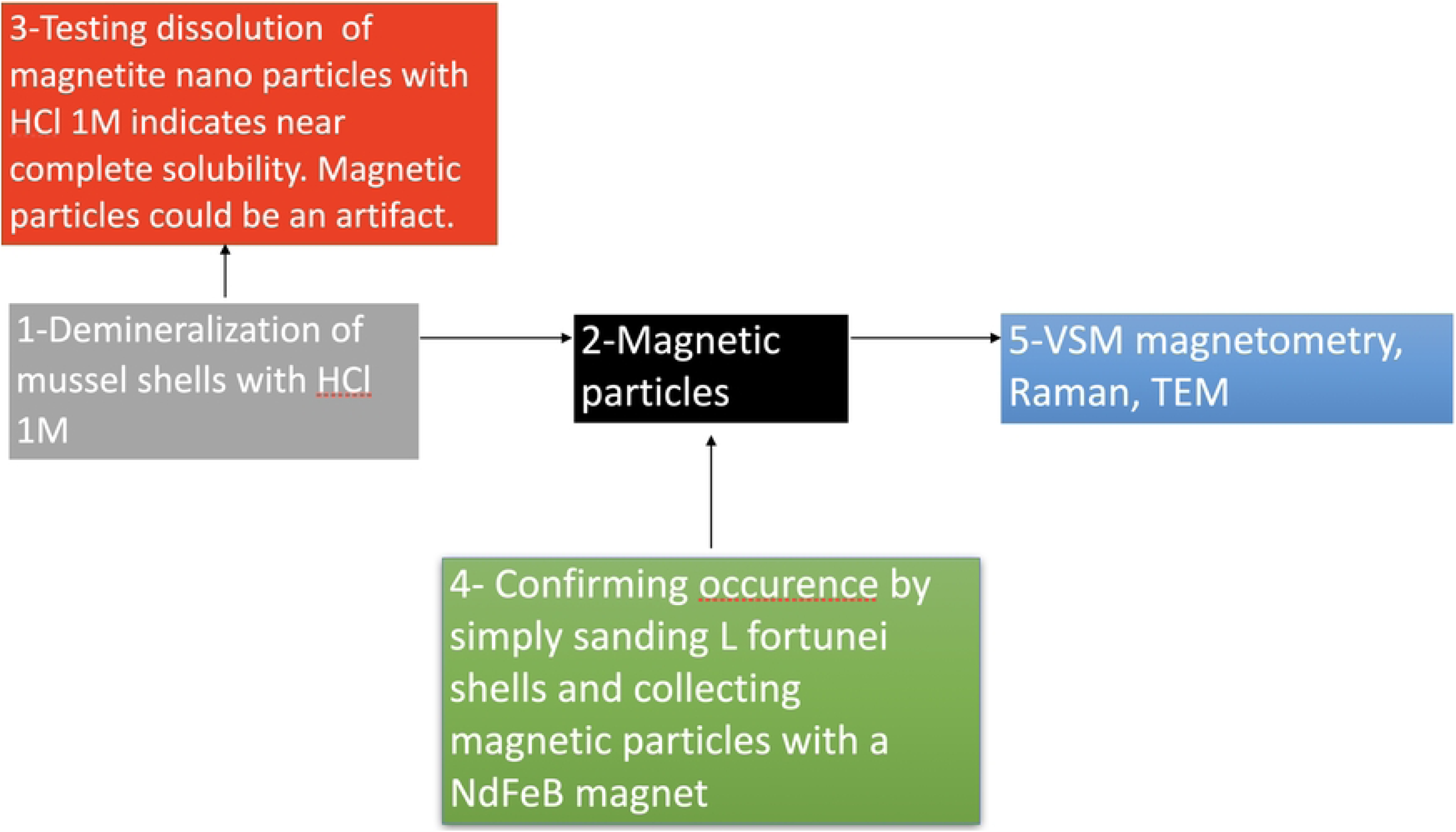
Diagram of the experimental procedure to verify if the magnetic particles were an artifact of shell demineralization with HCl 1M. Just a simply sanding with commercial sandpaper were enough to collect magnetic particles.

The role of the magnetic particle and their location in the shell are still under investigation but TEM electron microscope experiments presented here show that they are nanoparticles. Coupled with VSM magnetometer results, that show a faint ferromagnetic signal throughout the shells and indicate that they are well distributed and not concentrated. With individual particles with 10 nm diameter they might be homogeneously distributed through the entire volume of the nacre. This emphasizes their play on the self organization process of the shell construction, more than a possible magnetic orientation function. If this happens to be close to the actual shell construction process then the self organization, at least on the bivalve shell construction, are helped by ferromagnetic forces.

Other experiments presented here reinforce this observation as a simple grinding of the shells with comercial sandpaper (SiC, silicon carbide grinding media) produces collectable magnetic particles from the shell of bivalves. Anyone with a bivalve shell (maritime or freshwater) and sandpaper can replicate the results (See Supporting S1- Video: Sanded L fortunei mussel shell: collecting magnetic particles. Cleaned and dried Limnoperna fortunei shell were sanded with commercial sandpaper (SiC)).

Finally, both theories (protein as main actors and inorganic carbonate as main actors) agree on one point that the self organizing carbonate aggregates are amorphous and turn to crystalline as they gather at the growing aragonite site. As the transition from amorphous to crystalline is exothermic these processes burde no energy cost for organisms.

## Acknowledgments

Our thanks go to CEMIG S.A (www.cemig.com.br), Minas Gerais, Brazil, for the financial support (Project GT 604 – Control of the golden mussel: bioengineering and new materials for application in ecosystems and hydropower plants, www.cbeih.org).

## Supporting Information

S1- Video: Sanded L fortunei mussel shell: collecting magnetic particles. Cleaned and dried Limnoperna fortunei shell were sanded with commercial sandpaper (SiC).

S2- Video: Collecting magnetic particles from demineralized Perna perna mussel shell

S3 Table - Energy Dispersive X-Ray Fluorescence (EDX) results (in ppm) of the atomic elements present on shell of the freshwater Limnoperna fortunei bivalve and on the marine P perna Bivalve. Equipment: EDX 7000 Shimadzu.

